# Inherited flagellar structures coordinate the onset of swimming among related *E. coli* cells

**DOI:** 10.64898/2026.09.21.753120

**Authors:** Divya Choudhary, Youlian Goulev, Noah Olsman, Carlos Sanchez, Johannes M. Keegstra, Ethan Garner, Quincey Justman, Philippe Cluzel, Johan Paulsson

## Abstract

Bacteria commit to costly behaviors before they can benefit from them. In *Escherichia coli*, flagellar synthesis is expensive, yet the genes are expressed in stochastic pulses and only a small minority of pulses result in motility. Despite this noise, we find that lineage-related cells begin swimming at similar times, in groups of two to eight cells. Rare, high-amplitude pulses commit a single cell within one generation, but more commonly, pulses rise slowly and fail to trigger swimming before division. Rather than decaying, their output accumulates as cells inherit hook-basal bodies (HBBs) and continue to build new ones. This structural inheritance acts as a physical integrator, summing ongoing transcriptional activity across generations, so that an incomplete flagellar cascade is carried forward at division rather than lost. HBB accumulation sets when cells swim, and the anti-sigma factor FlgM sets how sharply. By holding late flagellar genes off until several HBBs have accumulated, FlgM makes motility depend steeply on early flagellar gene expression, so that relatives inheriting similar sets of HBBs become motile together. Without FlgM, this sharp dependence is largely lost, and activations of the full cascade do not reliably produce swimming. Together, our results show how the accumulation of a molecular machine integrates fluctuations, and how a checkpoint on its assembly sharpens them into a decision, setting both the speed and heritability of a behavioral trait.

## INTRODUCTION

Bacteria rarely experience environments that remain favorable forever. Nutrient or chemical gradients in the gut, soil or on surfaces change over hours or days, forcing cells to balance the benefits of remaining in one patch against the cost of searching for a new one. Facing this trade-off, genetically identical cells can diversify, by exploiting noisy gene expression^1^ of regulatory genes to randomize their swimming behaviors. Chemotaxis has been a model system for understanding such phenotypic variation, explaining how differences in the exact protein composition across cells bias movement along the same gradients^2–6^. How a cell commits to motility in the first place, transitioning from a state in which no flagellar genes are expressed to one where it starts swimming, is much less understood^7^.

Molecularly, flagellar synthesis is governed by a three-tiered cascade^8^. The master regulator, FlhDC (Class I), activates ∼30 genes (Class II) that in turn direct the assembly of the hook-basal-body (HBB), which anchors the filament structure to the cell envelope^9–11^. FlhDC also promotes expression of the alternate sigma factor FliA, which co-activates later components of HBB formation^10^. To prevent premature filament construction, the anti-sigma factor FlgM binds FliA until the HBB is fully assembled and able to secrete FlgM out of the cell^8,12–14^. Only then does FliA activate the ∼20 genes (Class III) that build the filament^15–18^.

However, quantitative assays for understanding the dynamics of this cascade^18,19^ have largely relied on population-average measurements or experiments that track already swimming cells^20–24^, often while using strains where the natural control circuits have been circumvented to overexpress the master regulator of flagellar synthesis, whether constitutively-motile mutants^25–28^ or inducible flagellar synthesis circuits^13,15–17,29,30^. Most individual cells then behave alike, expressing flagellar components at high levels such that transitions from induction to motility occur in most cells and during a single cell cycle^13,30^.

Such experiments can reveal the kinetics of flagellar assembly, but wild-type *E. coli* instead express flagellar genes at very low average levels, in rare and highly variable pulses^7,12,29,31^ that can last several generations^7^. Studies of non-motile *E. coli* further show that some pulses of Class II are not accompanied by a Class III pulse^7,12^, suggesting that not all cells that initiate the cascade become motile. This raises several questions. Given that a cell must invest in motility before it can use it, how far ahead does it commit? And does one individual cell commit to motility per initiation event, and then swims out as an isolated scout while its relatives remain in place, or does that commitment persist for generations to send a whole lineage of sister or cousin cells out in search of new niches? Furthermore, building flagella^22,32^ is a major undertaking that consumes several percent of the total cellular protein budget^33–35^ and requires coordinated expression of dozens of genes^8^ and subsequent assembly of gene products^7,9,17,18,24,36^. Given that wild type *E. coli* cells only rarely, transiently and weakly express these systems, how do they avoid making partially finished flagella that impose much of the burden but provide none of the function?

Here we address these questions by tracking the multigenerational transcriptional history that precedes motility while also monitoring motility itself, linking stochastic expression events within individual bacteria to the collective dispersal strategies^37–39^. Specifically, we follow Class II and Class III expression in wild type cells as well as in constitutively motile and non-motile control strains and use modified mother-machine devices^40^ to track multigenerational behaviors, including the onset of swimming. We find that for our wild type strain, Class II pulses occur in 15% of cell generations, and that fewer than 5% of those pulses are followed by transient Class III activation and swimming, typically two to three generations after Class II activation. These rare transitions frequently involve multiple relatives – sisters, cousins and second cousins – that all activate Class III and become motile within a narrow time window. A single ancestral Class II pulse thus produces several swimmers. This lineage coordination arises through the gradual accumulation and inheritance of HBBs assembled from Class II products. When HBBs accumulate rapidly over time, it produces motility within a single generation, whereas slowly rising Class II expression accumulates HBBs across divisions. Completed HBBs then enable FlgM secretion^8,12–14^, releasing FliA to activate Class III genes and coupling inherited structural progress to motility onset. If assembly restarted at each division, weak Class II pulses would incur the cost of building HBBs without ever delivering motility. Structural inheritance removes this penalty. Descendant cells build on what their ancestors made, so synthesis in a cell that never swims will still support swimming generations later. FlgM ensures that inherited HBBs do not trigger the late program prematurely. HBB accumulation integrates Class II expression across generations and sets when cells become motile, while FlgM sets how sharply they do so. Class III genes activate only once HBBs have accumulated to export FlgM, so cells that inherit HBBs but have not yet accumulated enough to swim within that generation keep Class III genes off. Because motility depends sharply on Class II expression, relatives that inherit similar sets of HBBs become motile at nearly the same time, activating Class III and swimming within a narrow window. Without FlgM, motility no longer depends sharply on Class II expression, and most activations of the full cascade fail to produce swimming. Thus, the flagellar cascade preserves incomplete assembly across generations, converting stochastic transcription in individual cells into coordinated dispersal across a lineage, without cell–cell communication.

## RESULTS

### Direct observation of *E. coli* flagellar cascade during transition from non-motile to motile

To monitor how an individual cell transitions from a non-motile to a motile state, and which transcriptional events precede that transition, we use modified versions of the mother-machine – a microfluidic device consisting of narrow, dead-end growth channels that retain a single “mother” cell while allowing daughter cells to be displaced by continuous medium flow, enabling long-term time-lapse imaging of individual-cell growth and behavior **[Figure 1A]**. Because cells in rich growth conditions have motility rates below our detection limit, and little need to swim, we continuously run medium that was first conditioned by prior growth (hereafter called conditioned medium, Supplementary Information). **[Figure 1B, S1]**.

**Figure 1.**
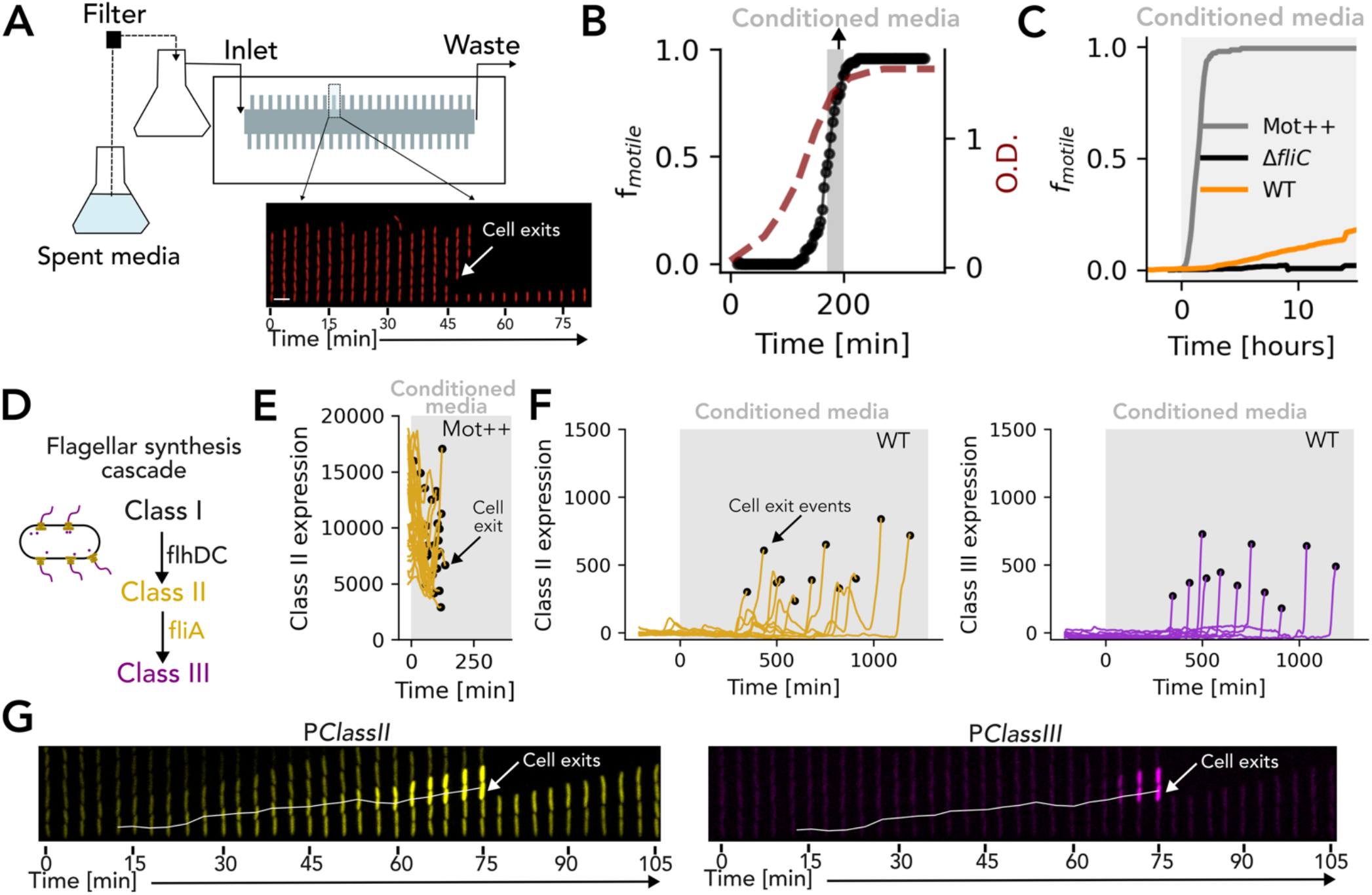
Visualizing gene expression leading to motility decision at the single cell level. (a) Schematic of the microfluidic experimental setup. Conditioned medium is supplied via an inlet and flows across perpendicular growth channels (trenches) where cells are imaged. The diagram illustrates a motile cell (marked by arrow) actively swimming out of the trench over time. (b) Temporal dynamics of the fraction of motile cells (black), and bulk culture optical density (dashed maroon line). The hyper-motile insertion mutant (Mot++ strain) exhibits motility during late exponential growth (conditioned media, marked in grey shaded region). (c) Cumulative fraction of motile cells (f_motile_) tracked after fresh media was replaced with conditioned media at t = 0 hours for WT (orange), Mot++ (grey) and a non-motile Δ*fliC* mutant (black). (d) Schematic of the hierarchical flagellar synthesis regulatory cascade, progressing from the master regulator (Class I, FlhDC) to structural genes and the FliA sigma factor (Class II), and finally to the motor and filament components (Class III). (e) Single-cell traces of Class II expression in motile Mot++ mother cells. The grey shaded region denotes the switch to conditioned media. Black dots mark the moment of cell exit (motility onset). (f) Single-cell traces of (Left, yellow) Class II and (Right, purple) Class III expression in motile WT mother cells. The grey shaded region denotes the switch to conditioned media. Black dots mark the moment of cell exit (motility onset). (g) Representative kymographs showing (left) Class II (*PClassII*-mVenus, yellow) and (right) Class III (*PClassIII*-mVenus, purple) promoter activity in WT cells. White arrows mark motility onset.

The confines of the mother machine may constrain actual swimming, but here we only track its onset, using exit from the growth channel as the readout. To confirm that the platform is accurate for that purpose, we switched from rich to conditioned growth medium and tracked both a non-motile strain lacking flagellin (Δ*fliC*) and a constitutively motile strain (hereafter Mot++, selected in our WT background, see Supplementary Information). All Mot++ cells exited within 1.40 ± 0.67 hours, as expected from the interval observed between flagellar cascade induction and movement^29,30,35^ **[Figure 1C-E, Movie 1]** and the motile cells could be directly observed swimming back and forth within the channel before exiting **[Movie 1-3, S2A].** The non-motile Δ*fliC* cells by contrast showed no such movement or exits **[Figure 1C]**. For WT we observed both behaviors, but exits were rare at approximately 0.5% per generation, and every exit was preceded by a rise in Class III expression based on a transcriptional reporter for FliA dependent transcription **[Figure 1C,1F-G, S2]**. Class III activation within a lineage reliably gave rise to motile progeny, with at least one motile cell produced in every case. Finally, to test whether the specific confinements of the mother machine impact how often cells commit to motility, we built an alternative microfluidic system where cells are held in place by flow-focus, using multiple small perfusion holes that hold swimming cells in place using opposing flow **[Figure 2A, Movie 4]**. The opposing current of growth medium is now too strong to overtake by swimming, and even motile cells are held in place. Using Class III expression bursts as a proxy for swimming reveals the same motility frequency as before, of approximately 0.5% per generation, matching the estimate based on physical exit **[Figure 2B, S3]**.

**Figure 2.**
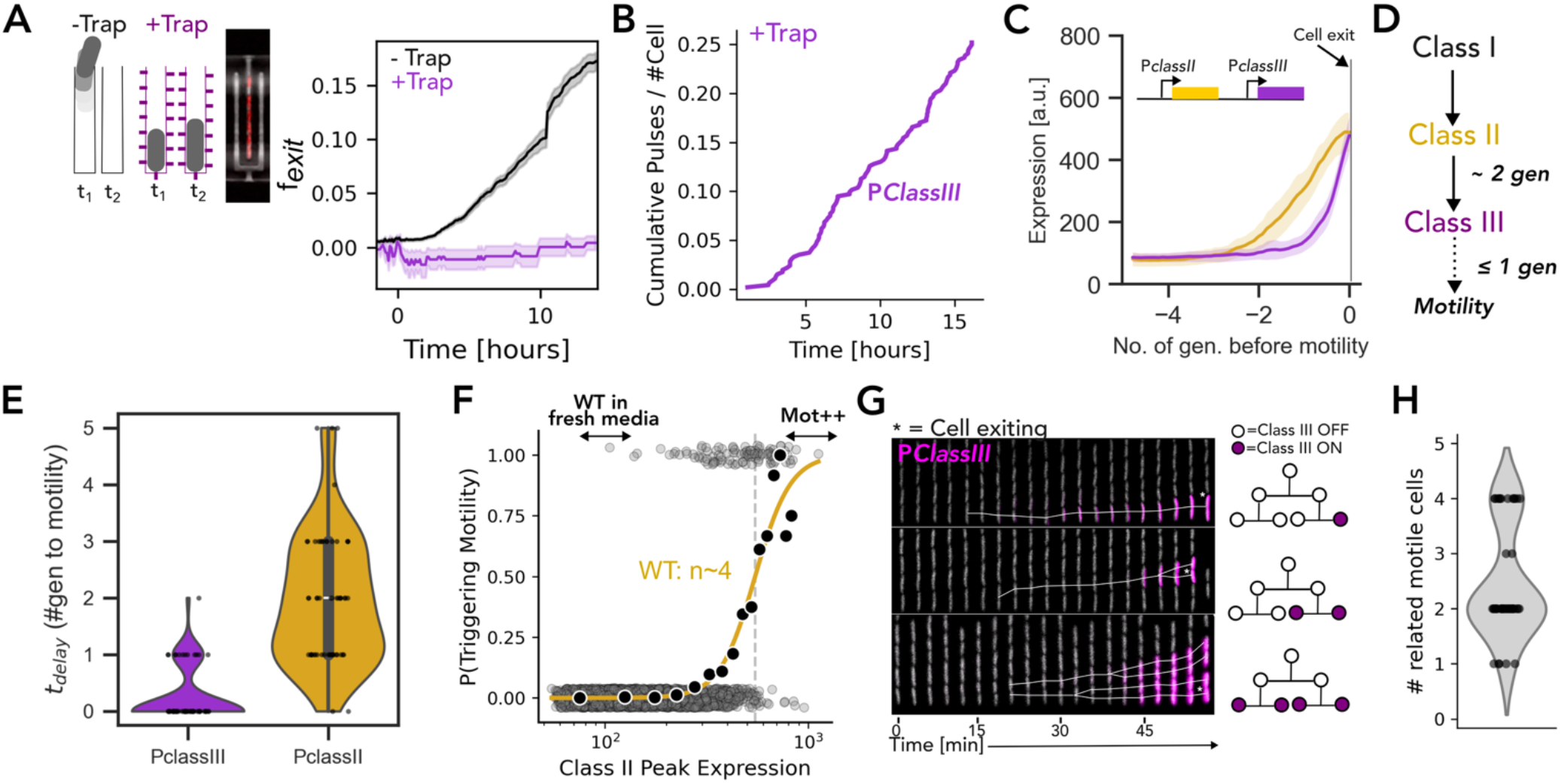
*E. coli* coordinate motility onset among relatives. (a) (Left) Schematic comparing the conventional microfluidic mother-machine (-Trap) with a modified design utilizing back-pressure to physically retain swimming cells (+Trap). (Right) Mean fraction of cells exiting the channel (f_exit_) after the provision of conditioned media at t = 0 for chips without (black) and with (purple) traps. Shaded regions represent standard deviation. (b) Cumulative number of Class III pulses per cell over time within the trap device. (c) Mean expression dynamics of Class II (*PClassIIsCFP3*, yellow) and Class III (*PClassIIImVenus*, purple) aligned to the time of motility onset (t = 0). Inset: Schematic of the dual-transcriptional reporter construct. (d) Schematic of the hierarchical flagellar cascade, denoting the temporal delays (∼2 generations for Class II, ≤ 1 generation for Class III) prior to motility. (e) Violin plots quantifying the delay (t_delay_, in number of generations) from the initial activation of Class II (yellow) and Class III (purple) promoters to the physical onset of motility. (f) Probability of triggering motility as a function of single-cell Class II peak expression. Grey scatter points represent individual Class II expression peaks, solid black dots indicate binned data points, and the yellow curve is a Hill equation fit. The fit reveals an apparent Hill coefficient of n∼4. (g) (Left) Kymographs tracking motile cells (marked by asterisk) just before they exit the growth trenches. Individual cell lineages are colored by Class III expression levels. The kymographs display activation of Class III in 1 (top), 2 (middle) and 4 (bottom) related cells. (Right) Lineage tree across two generations depicting the scenarios shown in left. Colored circles represent cells activating Class III expression. (h) Violin plot quantifying the number of related motile cells (descendants within an individual trench).

With the controls and basic measurements above in place, we next ask which regulatory events distinguish the rare WT cells that become motile. We built dual reporters for Class II and Class III transcription on a low-copy plasmid and introduced them into both WT and Mot++ strains, tracking cells that exit the channels alongside those that remain. Mot++ cells show high, relatively uniform Class II expression, and after the switch to conditioned medium rapidly upregulate the flagellar expression and exit **[Figure 1E, S4, Movie 1]**. WT cells, by contrast, show lower and more heterogeneous Class II expression **[Figure 1F, S5A-B]**. In fresh medium, WT Class II pulses are weak, and neither Class III activation nor channel exit is detected **[Figure 1F, S1, S5B]**. Switching to conditioned medium increases both the frequency and the amplitude of these pulses, to approximately 0.15 pulses per cell per generation **[Figure S5C, Movie 2]**. Only 3.3 ± 0.1% of pulses are followed by Class III activation, which quantitatively accounts for the approximately 0.5% of WT cells that become motile per generation **[Figure S5D]**. Most Class II pulses therefore end without completing the cascade, yet once Class III is activated, cells reliably become motile. The decision to swim is thus made between Class II and Class III activation, which we examine next.

### Completion of the flagellar cascade is delayed by generations after Class II activation

Tracking Class II and Class III expression over multiple generations in motile lineages reveals that the two classes do not necessarily activate within the same generation **[Figure 1G, 2C, S6A-B]**. In fact, Class II activation precedes Class III activation by an average of 1.74 ± 0.9 generations, while Class III appears as a sharp burst immediately before motility onset, within one generation of channel exit (0.25± 0.47 generations) **Figure 2C-E, S6A-B, S2]**. Because Class II genes encode the components required for hook–basal-body (HBB) assembly, we ask whether stronger Class II pulses are more likely to be followed by a Class III pulse and motility. Both the amplitude and duration of these pulses vary widely between cells **[Figure 1F, 2F, S5A-B]**, but contrary to our naïve expectation, larger Class II expression pulses did not yield a proportional increase in the probability of becoming motile. Instead, this probability remains low across a wide range of weak pulses before rising sharply over a narrow range of expression **[Figure 2F, S6C-D]**. This steep dependence is well described by a Hill function with an apparent Hill coefficient of approximately 4, revealing a highly nonlinear and apparently cooperative relationship between Class II expression and commitment to swimming. Motility therefore emerges as a threshold-like response to Class II activation. Mot++ cells predominantly express Class II above this threshold and enter motility nearly population-wide, whereas WT cells are predominantly subthreshold, with only rare cells crossing the threshold **[Figure 1E, 2F**].

The multi-generation delay following the rare Class II pulses could allow the initiating cell to give rise to one or more motile descendants. We therefore followed the timing of Class III activation among the relatives of each motile cell and asked how an ancestral Class II activation resolves motility within the growing lineage **[Figure 2G]**. We find that motility rarely arises as an isolated single-cell event. A Class II activation in a cell that eventually becomes motile is instead frequently propagated across divisions **[Figure 2H, S7, Movie 3-4]**. The tightest coordination is between sister cells, but we observe coordinated pulses of Class III for up to four cousin descendants from a shared grandmother, all occurring within a narrow time window **[Figure 2H, S2, S7A]**. A single founding Class II activation therefore commits several cells, rather than the one expected if commitment had been confined to the initiating generation. Given that a Class III pulse is rare, occurring roughly once every 100 generations, the odds of two cousins independently firing in the same time window are very small, and four to eight relatives doing so would not happen by chance at any measurable frequency. The decision to swim is thus inherited across a lineage, with information from a transient Class II activation propagating through successive divisions before being converted into coordinated motility. Two features of the mother machine complicate this measurement. First, cells leave through the open end of the channel, so a cell exiting from a deeper position displaces those above it, and the departure of those upper cells cannot be attributed to their own motility. To separate the two, we identify lineages where, by chance, the cell nearest the opening leave first and a deeper cell followed shortly after. Within these cases, that are easier to interpret, we found that each exit was preceded by its own Class III pulse, which begins shortly before swimming, confirming that clustered departures reflect genuine, coordinated motility **[Figure S2]**. Second, exit from the channels truncates a motile lineage at the moment it commits, so the full extent of coordination cannot be measured in the mother machine. We therefore return to the device with perfusion holes and flow-focus, which retains the swimming cells **[Figure 2A, S7]**. With all cells retained, motility – based on Class III expression pulses – again resolves as a coordinated multi-generational decision, confirming that the pattern persists when motile descendants remain observable **[Figure S7, Movie 4]**. We also observed groups of up to eight related descendants sharing a great-grandmother committing to motility within a narrow time window **[Figure S7B-C]**. This commitment is transient, where lineages maintain Class III expression for one to two generations **[Figure S7D]**. Coordination therefore amplifies the output of a rare and transient activation, yielding several motile descendants.

### Slow basal body accumulation drives simultaneous motility onset across lineages

Next, we address how a transcriptional pulse, initiating in a single ancestor, gives rise to near simultaneous Class III activation several divisions later. In principle, a shared extracellular signal like quorum sensing^41,42^, could synchronize nearby cells and also explain why a whole lineage of descendants activates swimming collectively. Such a signal, however, would be expected to couple cells by proximity rather than ancestry. Instead, we observe coordination along a family tree. Sisters and cousins fire Class III together, while unrelated neighbors in the same growth channel do not. Hence, the coupling mechanism must be inherited rather than signaled **[Figure 2, S7]**. Intuitively, multi-generational coordination can stem from the passive dilution^43^ of a large initial pool of Class II products, where a burst of Class II expression pulses dilutes through many rounds of divisions such that progenies have enough Class II products to become motile implying a larger delay **[Figure 3A-B]**. Such dilution has been observed for coordinated decisions to become motile in *Bacillus subtilis*^44^. However, our single-cell tracking of Class II expression from activation to motility onset reveals the exact opposite behavior **[Figure 3C]**. Cells with a larger burst of Class II pulses commit to motility faster, whereas lower Class II expression rates resulted in longer, multi-generational delays **[Figure 3D]**. This relationship reveals that cells are not merely diluting a transient repressor signal but rather accumulate Class II products across generations **[Figure 3E]**. Under this integrative framework, commitment timing is dictated not by the total amount of Class II products, but by the rate of Class II expression **[Figure 3C-D]**. This leads to delay in onset of motility inversely proportional to Class II activation rates.

**Figure 3.**
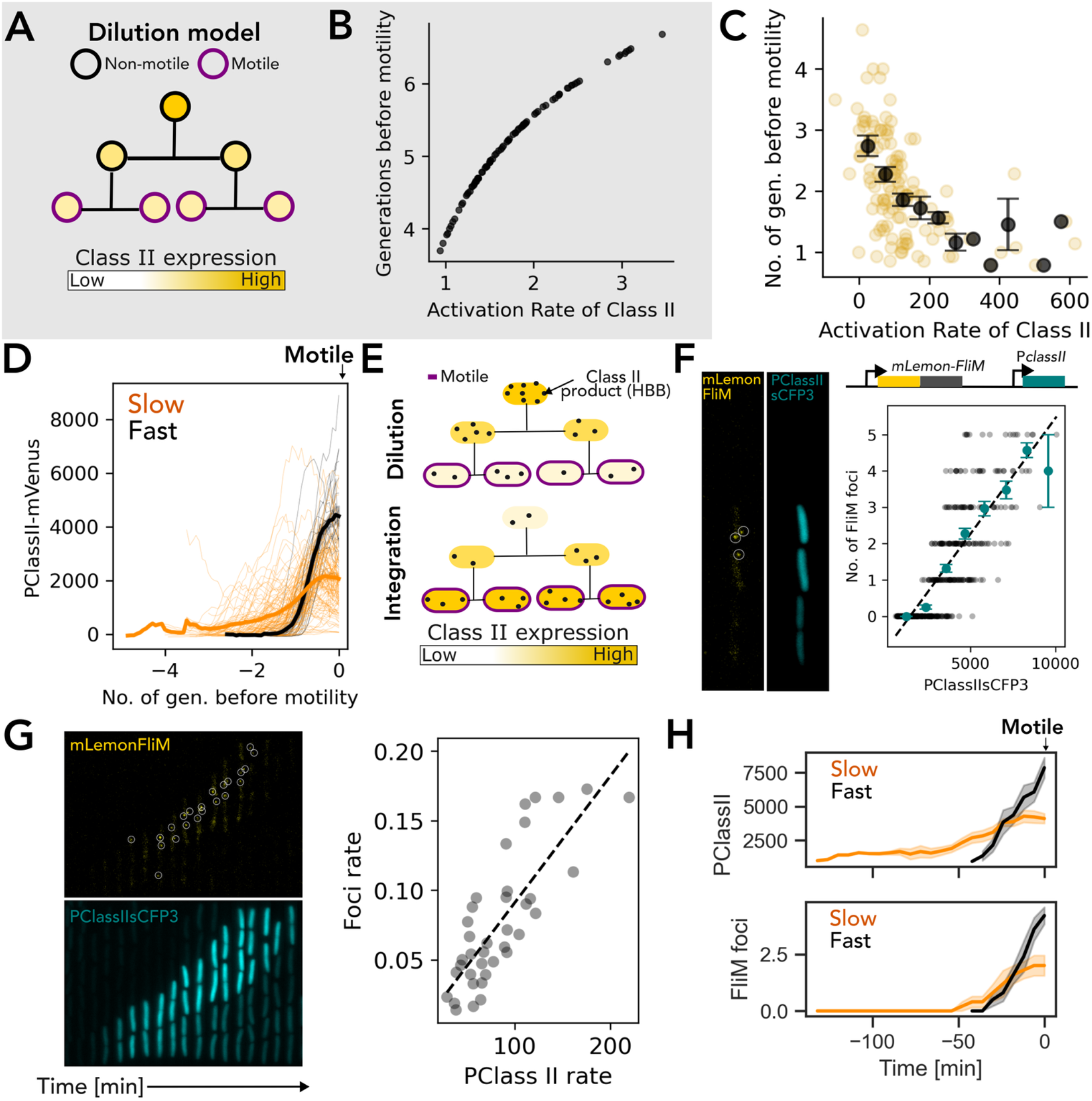
Rate of Class II expression and accumulation of HBBs governs coordination in motility onset. (a) Representative family tree for Class II expression across two divisions for the dilution model to understand coordinated onset of motility (motile cells outlined in purple, color of dots depicts expression of Class II). (b) Quantification of delay in motility behavior against the rate of Class II activity, where each colored scatter is an individual simulation. (c) Scatter plot of the number of generations to motility post Class II expression versus the rate of Class II expression. Each scatter point represents a motile cell, and black points show averages over bins of Class II expression rate. (d) *PClassII-mVenus* expression trajectories of motile mother cells aligned to the onset of motility (t = 0) for cells with fast (black) versus slow (orange) Class II transcription rates (average shown as bold lines). (e) Representative family tree for Class II expression (shown in yellow) and its products i.e. HBB (depicted by black dots) contrasting the coordination across lineage for the dilution (top) and integration (bottom) model. (f) (Left) Representative fluorescence snapshots and schematic of a dual-reporter strain tracking structural basal bodies (*mLemon-FliM*, yellow foci circled, left) alongside Class II transcription (*PClassII-sCFP3*, cyan, right). (Right) Single-cell correlation between the number of FliM foci and Class II expression intensity. Best linear fit marked with dashed lines (teal markers indicate binned means ± standard deviation). (g) (left) Kymographs tracking the intergenerational accumulation of HBBs (*mLemon-FliM*, top, white circles) and Class II transcription (*PClassII-sCFP3*, bottom) in a lineage. (Right) Scatter plot of the rate of FliM foci appearance with the rate of Class II transcription prior to motility. Each scatter point represents a motile cell. (h) Mean traces of Class II expression (top) and FliM foci accumulation (bottom) for fast (black) and slow (orange) Class II activity lineages.

To further test the predictions that multigenerational delays and coordinated motility are governed by the physical buildup of HBB components encoded by Class II, we next link transcriptional dynamics to visualizing HBBs. To do so, we combine a *PClassIIsCFP3* transcriptional reporter with a translational reporter for the C-ring protein FliM^45,46^ (*mLemonFliM*) **[Figure 3F, S8A-B, Movie 5-6]**. Dual-reporter measurements reveal a linear relationship between Class II expression and the number of *mLemonFliM* foci, demonstrating that transcriptional output is translated into HBB accumulation **[Figure 3F, Movie 6-7]**. Next, we follow the coupled dynamics in lineages that become motile **[Figure 3G]**. The rate of FliM focus accumulation closely tracks the rate of Class II expression **[Figure 3H]**. Rapid Class II activation produces a burst of *mLemonFliM* foci, whereas slow Class II activation result in the gradual accumulation of *mLemonFliM* foci over multiple generations. In these slowly assembling lineages, HBBs are inherited by daughters and continue to accumulate after division **[Figure 3H]**. This inheritance is essential. A cell that cannot accumulate enough HBBs to swim within one generation would otherwise lose its investment at division, forcing daughters to restart assembly, so slow Class II pulses would never produce swimmers. Instead, HBB foci partition between daughters rather than being degraded, and each generation builds on the structures its ancestors made. Investment that fails to produce motility in one cell is carried forward and completed in its descendants. Furthermore, cells containing one to three foci initially exhibit long and variable delays before motility, whereas cells possessing more than three *mLemonFliM* foci transition to motility rapidly (typically within ∼15–20 minutes) **[Figure S8A]**. Class III activation follows the same pattern. Dual reporters of Class III expression and *mLemonFliM* show that Class III activates after foci accumulation, rather than when the first focus appears **[Figure S8D-E, Movie 8]**. Cells therefore accumulate a set of basal bodies before activating Class III. HBB accumulation is thus the mechanistic basis of the delay separating transcriptional activation from behavioral output.

This structural inheritance explains why motility, despite being rare and transient, often emerges as coordinated bursts among related cells rather than isolated single cell events **[Figure 2, S7D]**. When HBB accumulation is slow relative to the cell cycle, partially assembled basal bodies are inherited across divisions, allowing multiple descendants to collectively integrate over the Class II activity and complete the flagellar program together **[Figure 3]**. In contrast, rapid HBB accumulation enables commitment within a single generation, producing isolated motility events. The rate of Class II expression therefore sets the yield of an activation as well as its timing. Slow accumulation converts one activation into many related swimmers, whereas rapid accumulation produces a single swimmer. In all, Class III activation rapidly triggers motility, and the multigenerational integration of Class II products acts as the primary timer preceding motility onset. This delay functions as a period of structural integration, transforming transient transcriptional fluctuations into a heritable lineage-level decision. Stochastic Class II activation thus not only seeds phenotypic variation, but via its rate of expression dictates the timing and lineage coordination of the behavior for progenies.

### FlgM sharpens the conversion of flagellar expression into motility

Above we showed that Class II and Class III transcription are separated by several generations in WT cells, and that Class III activates shortly before swimming begins. If HBB accumulation, rather than the transcriptional checkpoint, sets this delay, then releasing Class III early should not advance swimming. We test this prediction by removing the checkpoint separating the two classes. In the established flagellar cascade, the anti-sigma factor FlgM binds FliA and inhibits Class III transcription until HBBs become competent to export FlgM from the cell^8,12–14^ **[Figure 4A]**. Deleting *flgM* should therefore allow downstream transcription to begin without waiting for assembly. As expected, Class II and Class III activate together in *ΔflgM* cells **[Figure 4B, S9A]**. Yet motility remains delayed by several generations after Class II activation, as in wild type **[Figure 4B, S10, Movie 9]**. Activating both classes of genes together is therefore insufficient to produce immediate swimming, and cells must still accumulate the structures needed for motility.

**Figure 4.**
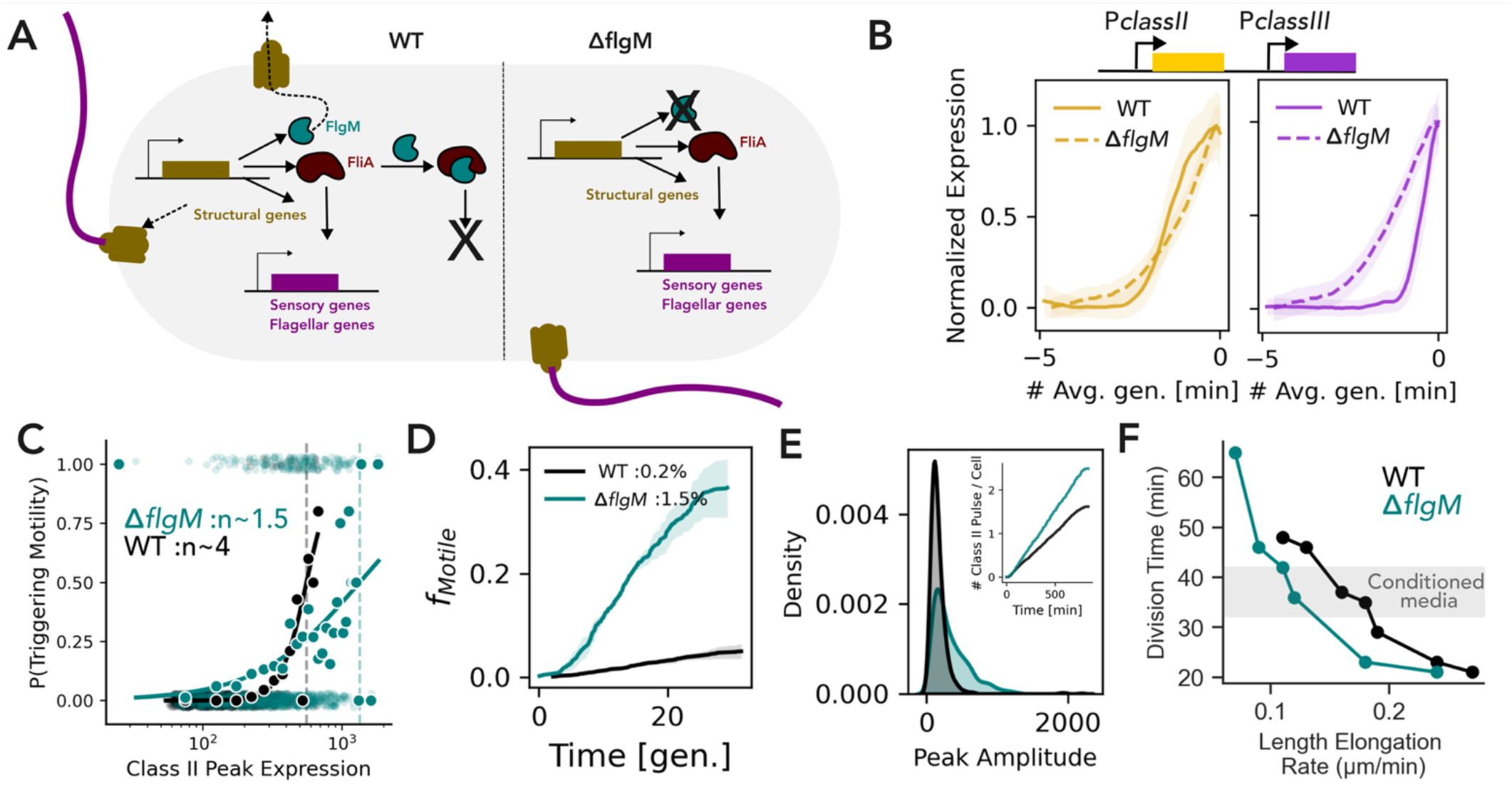
FlgM sharpens the conversion of flagellar gene expression into motility. (a) Schematic comparing the flagellar regulatory cascade in wild-type (WT) and *ΔflgM* cells. In WT, FlgM sequesters FliA to prevent premature Class III transcription until the hook-basal body is complete. In *ΔflgM*, FliA is prematurely liberated, uncoupling Class III activation from structural assembly. (b) Normalized average expression trajectories of Class II (yellow, left) and Class III (purple, right) genes aligned to the onset of motility (t = 0) for *ΔflgM* (dashed lines) and WT cells (solid lines). (c) Single-cell probability of triggering motility as a function of peak Class II expression for WT (black) and *ΔflgM* cells (teal) with estimation of apparent Hill coefficients. Light scatter points are individual data points for Class II peak expression, and the dark points are binned averages with the line plot showing the fit to estimate the apparent Hill coefficient. (d) Mean cumulative fraction of motile cells (f_Motile_) tracked over time for *ΔflgM* (teal) and WT (black) cells. Shaded regions represent standard deviation. (e) Probability density distribution of single-cell Class II peak amplitudes for *ΔflgM* (teal) and WT (black) cells. Inset: Cumulative number of Class II pulses per cell over time for *ΔflgM* (teal) and WT (black) cells. (f) Scatter plot of division time versus cell elongation rate across varying nutrient depletion conditions *for ΔflgM* (teal) and WT cells (black).

What, then, does the checkpoint contribute? In wild type, Class III activates only after several HBBs have accumulated, the stage at which FlgM is exported. We therefore asked whether removing this coupling alters how Class II expression is converted into motility. In wild type, only rare Class II pulses are accompanied by Class III activation, whereas in *ΔflgM*, nearly all Class II pulses activate Class III [**Figure 4B, S9B]**. The late program no longer waits for assembly. Motility begins to appear at a similar Class II expression level in both strains **[Figure 4C]**, but above this level the two strains behave differently. In wild type, the probability of swimming rises steeply over a narrow range of expression, whereas in *ΔflgM* it rises only gradually, and many cells remain non-motile despite reaching expression levels that almost always produce swimmers in wild type. The apparent Hill coefficient falls from approximately 4 to 1.5 **[Figure 4C]**. Without FlgM, the switch-like response becomes graded, because Class III activation no longer reads out how much a cell has assembled, and Class II expression therefore no longer reliably predicts whether a cell swims. Class III activation itself shows this directly. In wild type, Class III activation reliably predicts motility, because it occurs only after HBBs have accumulated to export FlgM. In *ΔflgM*, by contrast, more than 80% of Class III pulses are not followed by swimming **[Figure S9C, Movie 2, 9]**, as cells activate the late program before they have built enough HBBs to use it. Together, these changes undermine the basis for lineage coordination. In wild type, relatives that inherit similar sets of HBBs cross the same sharp threshold and become motile together. Without FlgM, this basis is lost in two ways. First, because Class III no longer reports HBB accumulation, sisters that inherit similar sets of HBBs need not activate the late program at the same time. Second, because the response is graded even at high expression, relatives with comparable Class II levels can still differ in whether they swim. Both effects would be expected to weaken the synchronous motility seen in wild-type lineages. HBB accumulation thus sets when cells become motile, while FlgM sets how sharply, so that Class II activation reliably predicts, and Class III activation reliably signals, impending swimming, allowing relatives that build similar sets of HBBs to become motile together.

Why does Class II expression become a poor predictor of motility without FlgM? The answer lies in the architecture of the fliA promoter, which responds to both Class II and Class III regulation and therefore allows FliA to upregulate transcription of its own operon^10,47^ [**Figure 4A**]. In wild type, this feedback is coupled to assembly, because FliA is released only as HBBs export FlgM, so that amplification of Class II expression tracks structural progress. In *ΔflgM*, the feedback no longer waits for assembly. Class II expression rises faster and reaches higher amplitudes **[Figure 4D–E, Movie 9, 10]**, but these levels no longer reflect how many HBBs a cell has built, so cells with similar expression can be at very different stages of assembly. The stronger expression still pushes more cells toward swimming. *ΔflgM* populations produce about seven-fold more swimmers than WT, and 15% of Class II pulses lead to motility, compared with 3% in wild type **[Figure 4D, S9]**, an increase driven by more expression rather than more effective expression. This additional swimming comes at a cost. Although mutant and wild-type cells have similar division times across the growth curve, *ΔflgM* cells elongate more slowly, most markedly at intermediate growth rates in conditioned medium, where motility is most frequent, consistent with a greater biosynthetic burden **[Figure 4F]**. FlgM therefore trades the number of swimmers for the reliability of those that do swim, balancing the expense of flagellar synthesis against the likelihood of realizing its benefit.

## DISCUSSION

Bacterial motility is a fundamental survival strategy ^48,49^, yet how individual cells time, execute, and inherit this decision has remained unresolved, largely because non-motile to motile transitions cannot be followed together with their gene-expression dynamics using standard experimental approaches. Bacterial decisions are often described as transcriptional thresholds that convert environmental signals into changes in behavior^7,50–52^. Our lineage-based observations extend that picture. Commitment to motility in *E. coli* depends not only on an instantaneous transcriptional switch, but on the kinetic race between stochastic gene expression, growth-dependent dilution, and the accumulation of hook-basal bodies. By mapping the flagellar cascade from initial promoter activation to physical channel exit, we find that transcriptional pulses of Class II genes are common and transient. Only 3% of these activations successfully lead to motility, establishing a low baseline motility rate of about 0.5% per generation. Crucially, transcriptional variability in Class II expression merely seeds heterogeneity. It is the rate of Class II expression and HBB accumulation relative to cell division that provides temporal integration and heritability. When HBB accumulation is slow relative to the division cycle, partially assembled HBBs are inherited across generations. This physical inheritance extends the integration window, effectively storing transient transcriptional memory in a structural intermediate and producing coordinated bursts of motility among related descendants. Conversely, when HBB accumulation proceeds rapidly, commitment occurs within a single generation, and the temporal coupling between transcription and movement tightens. The HBB therefore functions as a physical integrator, converting transient, noisy transcription into a coordinated behavioral state. Integration is paired with a sharp threshold. The probability of motility rises steeply with Class II expression (apparent Hill coefficient ∼4), so investment can accumulate across generations without producing swimmers and is converted into motility only once it crosses this threshold.

The physical inheritance of basal bodies is not only a timing mechanism but also changes the effective cost of motility. Commitment requires a cell to have accumulated basal bodies. This requirement for several complete structures, rather than one, is consistent with the steep dependence of motility on Class II expression and with the delay before Class III activation. If each cell had to reach the threshold autonomously within a single generation, subthreshold investments would be lost at division, imposing cost on the population without producing motility. Our results show that subthreshold investment is not necessarily wasted as the HBBs partition at division and accumulate across generations, allowing descendants to inherit and add to HBBs made by ancestors. Thus, the relevant cost of motility is not the total amount of flagellar material synthesized, but the fraction of that investment that never contributes to motility. Lineage coordination reduces this wasted investment by allowing HBBs to accumulate beyond a cell cycle. In this way, the lineage itself becomes a unit over which the cost of motility is distributed, as HBBs built by one cell can contribute to the motile phenotype of its descendants.

Our results also expand the role of the anti-sigma factor FlgM, classically regarded as a checkpoint that withholds filament synthesis until the hook-basal body is complete ^8,12–14^. In *ΔflgM* cells, motility remains delayed by several generations, and cells still accumulate multiple basal bodies before swimming, indicating that releasing the transcriptional checkpoint does not bypass the need for HBB accumulation. FlgM instead sharpens the conversion of this accumulated investment into behavior. Its export through completed HBBs couples Class III activation to assembly progress, helping make motility a steep, threshold-like function of Class II expression. Without FlgM, this coupling is weakened at both stages of the cascade. Class III activates irrespective of assembly, while FliA autoregulation can amplify Class II reporter expression without waiting for FlgM export, weakening its correspondence with basal body abundance. The sharpness conferred by FlgM may therefore contribute to lineage coordination. Related cells inherit basal bodies and continue to build on this shared investment. In wild type, Class III activation reads out this shared assembly, and the steep threshold ensures that relatives with similar Class II expression become motile together. Without FlgM, both links are lost. Because Class III is active regardless of how many basal bodies a cell carries, it can no longer report inherited structure, and because motility no longer depends sharply on Class II expression, relatives with similar Class II levels need not become motile, or may do so at different times. By coupling late-gene activation to inherited and newly assembled structures, FlgM thus helps translate a shared assembly history into a coordinated behavioral transition.

Finally, returning to the question we began with, the lineage scale of the decision sets how many swimmers a rare activation produces. Because commitment is inherited rather than signaled, cells leave as small groups of close relatives while the rest of the population stays behind and continues to hedge. These relatives would not stay together once they leave. Swimming trajectories decorrelate within tens of seconds^53^, so a lineage that departs as a group will not arrive as one. Lineage coordination therefore changes how many independent attempts a single activation produces. In environments where most cells never encounter a patch, the growth of the population depends on the rare cells that do, and the number sent out sets how likely it is that any one of them founds a new colony^6,54^. If establishment carries a threshold, as it does where colonization is cooperative, cells that did arrive together would establish more readily^55^, though our measurements do not resolve whether relatives remain close enough for this to apply. Inheritance can be more robust to conditions that disrupt signaling^56^. Under flow, or at low cell density, a secreted signal is diluted before it can accumulate^56^, whereas inheritance is not, so a lineage keeps converting to the motile state under conditions in which a diffusible cue would diffuse away.

The implications of this structural integration extend beyond flagellar motility. The *E. coli* flagellar export apparatus belongs to the broader superfamily of type III secretion systems (T3SS), which includes the injectisome ^8,28,57^, that is a structurally homologous needle complex utilized by pathogens such as *Salmonella* ^58^ and *Yersinia* ^59^ to deliver effectors into host cells. Both systems share a highly conserved export gate and a substrate-specificity switch that orders export^36,60^. Given this shared architecture, it is plausible that the assembly-dependent mechanisms we observe in *E. coli* also shape when, and in which cells, pathogens deploy their secretion systems. Structural inheritance may similarly contribute to the heritable expression of multiprotein structures, such as R-bodies^61^. Overall, the flagellar cascade exemplifies a form of bacterial decision-making in which the slow accumulation of molecular machines integrates transcriptional fluctuations, dictating not just the fate of a single cell, but the coordinated behavior of a lineage.

## Acknowledgments

We thank Maxence Vincent and members of the Paulsson, Cluzel and Garner labs for their discussions and comments on the manuscript. Research in the Paulsson lab is funded by ARPA-H. D.C. is supported by an HFSP postdoctoral fellowship.

## Author contributions

Conception and design of study, D.C.; engineering of genetic constructs, D.C. and N.O.; microfluidic setup, D.C. and C.S.; mother machine chip design and fabrication, D.C., and Y.G.; data analysis and interpretation, D.C., Q.J. and P.C.; writing and editing of the article, D.C., Q.J., J.P., E.G., J.M.K.

## Supplementary Information

**Figure S1.**
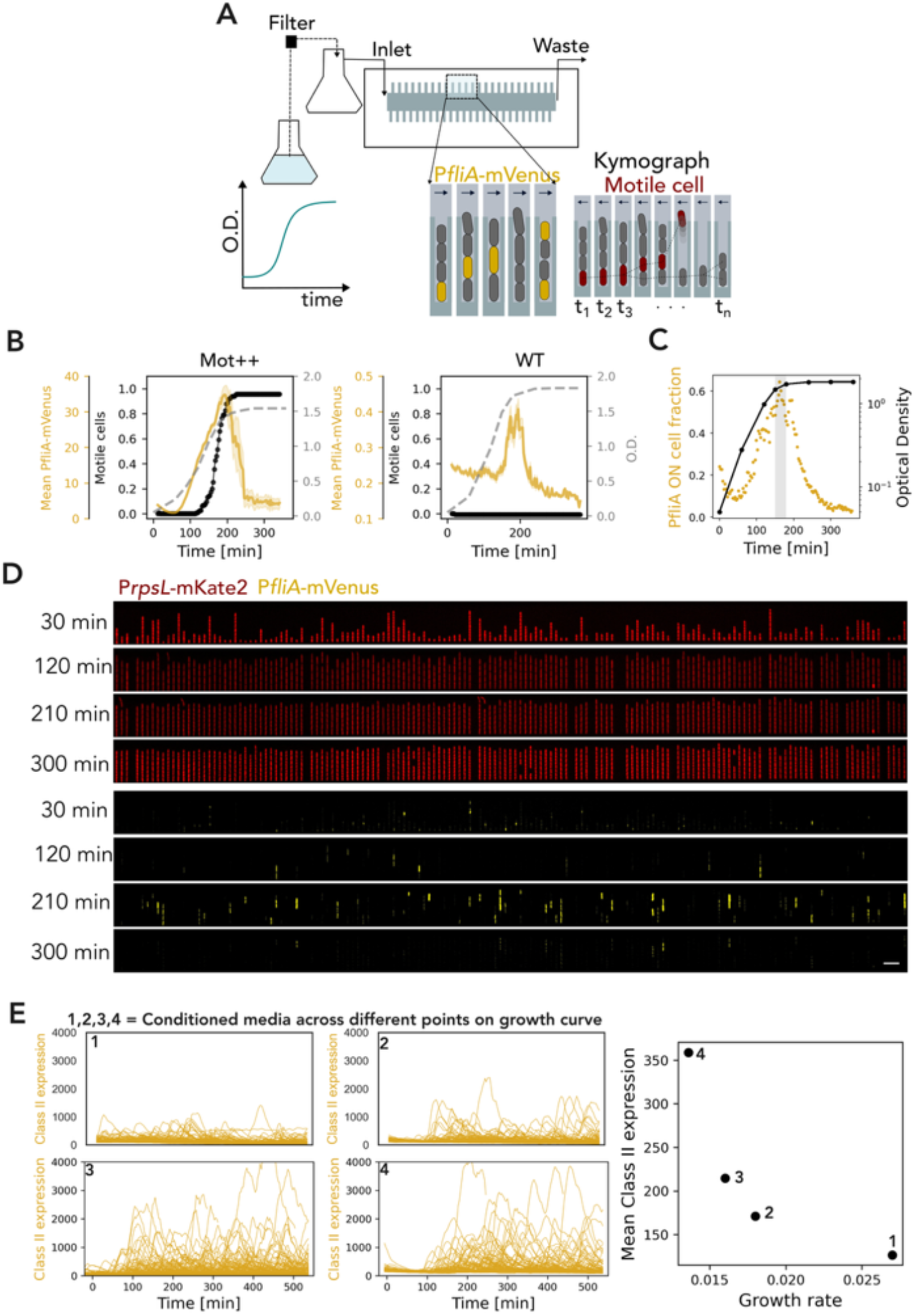
Population and single-cell dynamics of Class II flagellar gene expression across growth phases. (A) Schematic of the experimental setup. Media drawn from a bulk culture at specific optical densities (O.D.) is continuously flowed into the microfluidic mother-machine. The accompanying diagrams illustrate the tracking of Class II promoter activity (P*ClassII-mVenus*, yellow) and the physical exit of a motile cell (red) from the growth trench over sequential time points (t1 to tn). (B) Temporal dynamics of mean Class II expression (P*ClassII-mVenus*, yellow), the fraction of motile cells (black), and bulk culture optical density (dashed grey line) for (Left) the hyper-motile insertion mutant (Mot++ strain) and (Right) wild-type (WT) cells. (C) The fraction of WT cells actively expressing Class II genes (PfliA ON fraction, yellow) overlaid with the bulk culture growth curve (O.D., black). The Class II activation peaks during the mid-to-late exponential growth phase (grey shaded region). (D) Representative microfluidic image montages capturing WT cells at distinct temporal stages (30, 120, 210, and 300 min) along the growth curve for (Top) Constitutive marker (PrpsL-mKate2, red) and (Bottom) the Class II reporter (PfliA-mVenus, yellow). (E) (Left) Single-cell mother cell traces of Class II expression under four distinct steady-state growth conditions chosen as spent media from different points across the growth curve. (Right) Scatter plot of mean Class II expression versus the single-cell growth rate.

**Figure S2.**
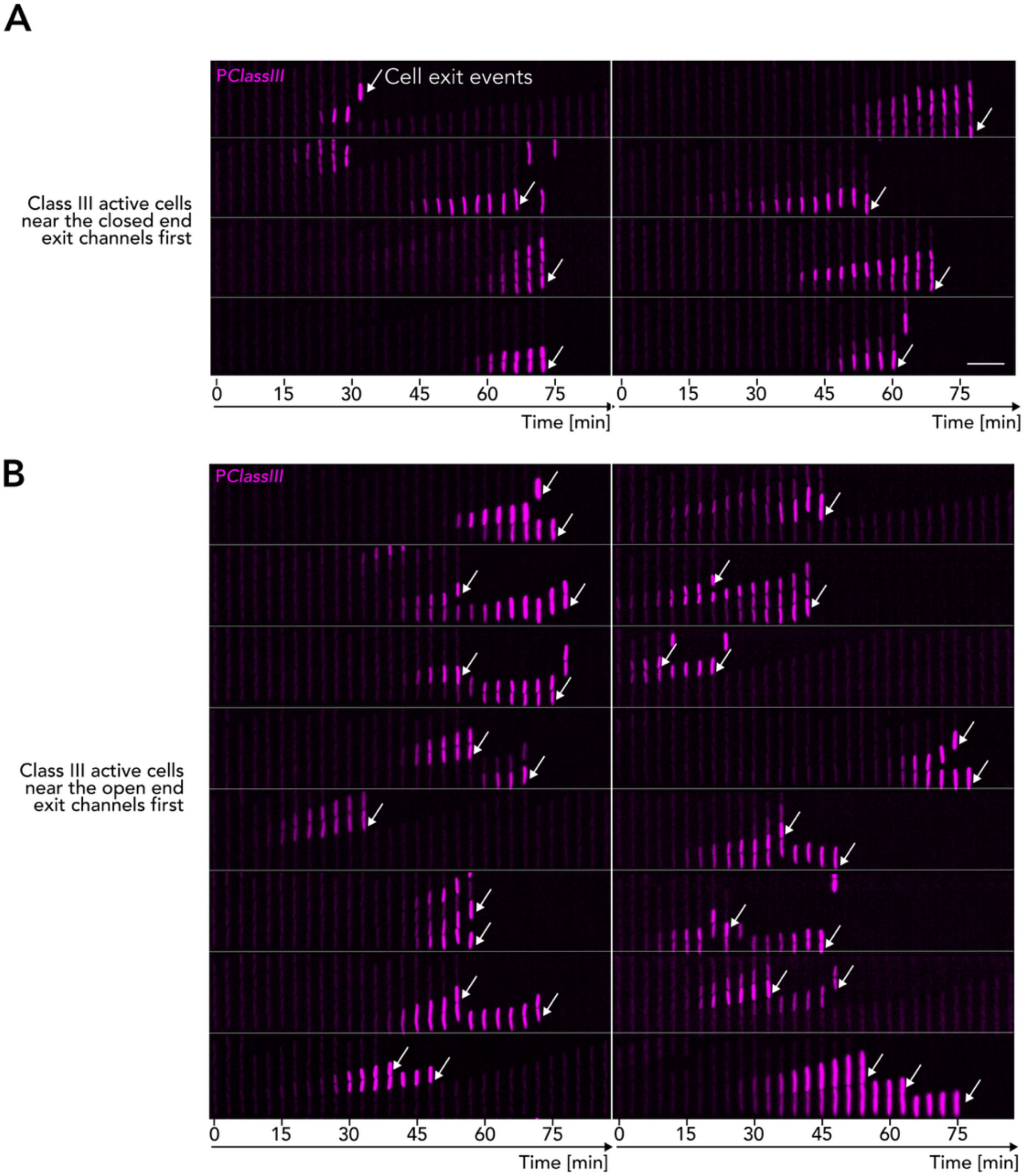
Coordinated class III activation before cell exit. Kymographs tracking cells (arrows) positioned (A) near the closed end or (B) near the open end of the growth trench, among coordinated Class III-active cells, just before they exit. Individual cell lineages are colored by Class III expression level.

**Figure S3.**
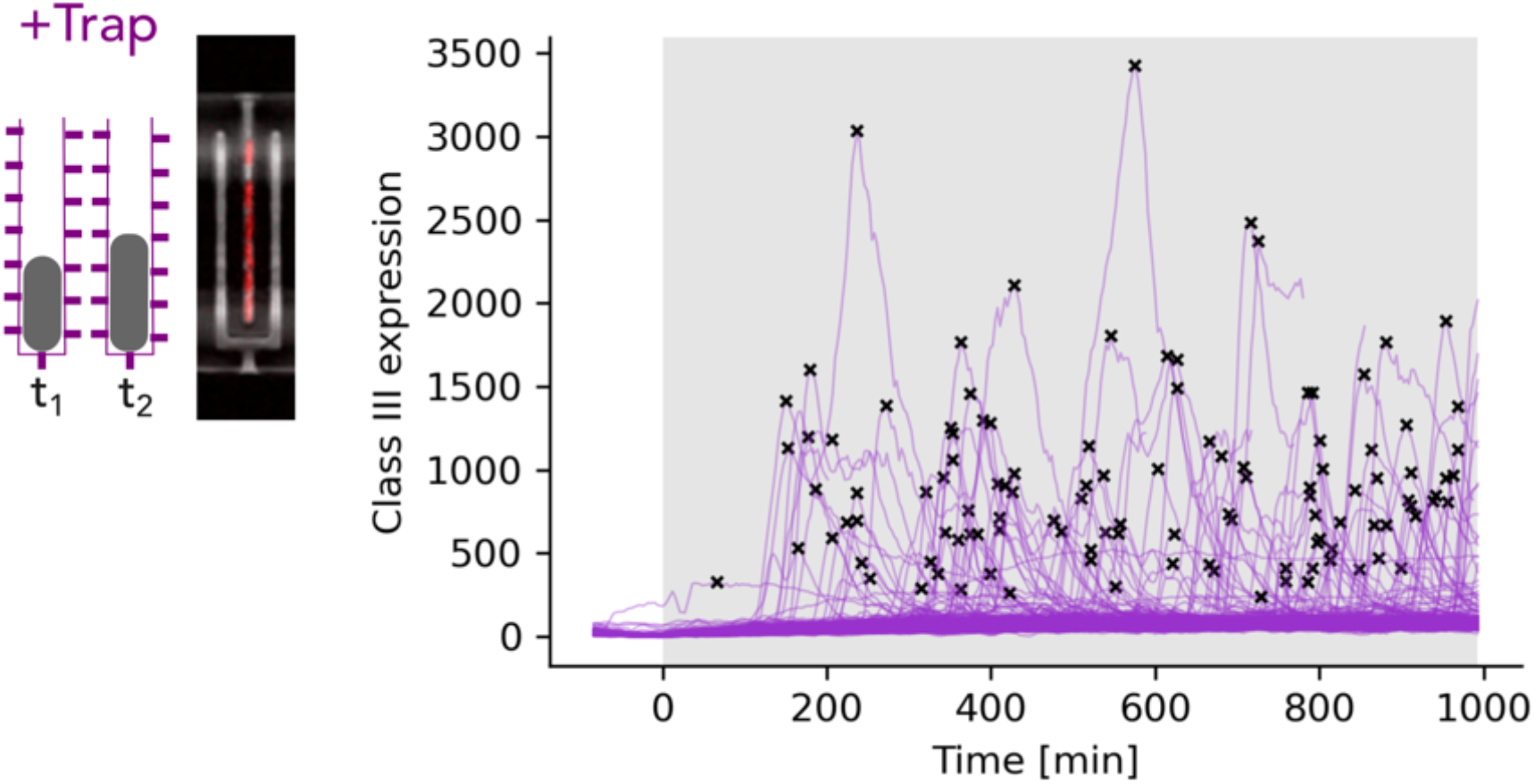
Microfluidic trapping preserves Class III transcriptional pulsing dynamics. (Left) Schematic comparing the conventional microfluidic mother-machine (-Trap) with a modified design utilizing back-pressure to physically retain swimming cells (+Trap). (Right) Single-cell mother cell expression traces of the Class III reporter (*PClassIIImVenus*) for lineages confined within the trap. Black crosses (’x’) mark detected transcriptional pulse peaks. The shaded grey region denotes the switch to conditioned media at t=0 mins.

**Figure S4.**
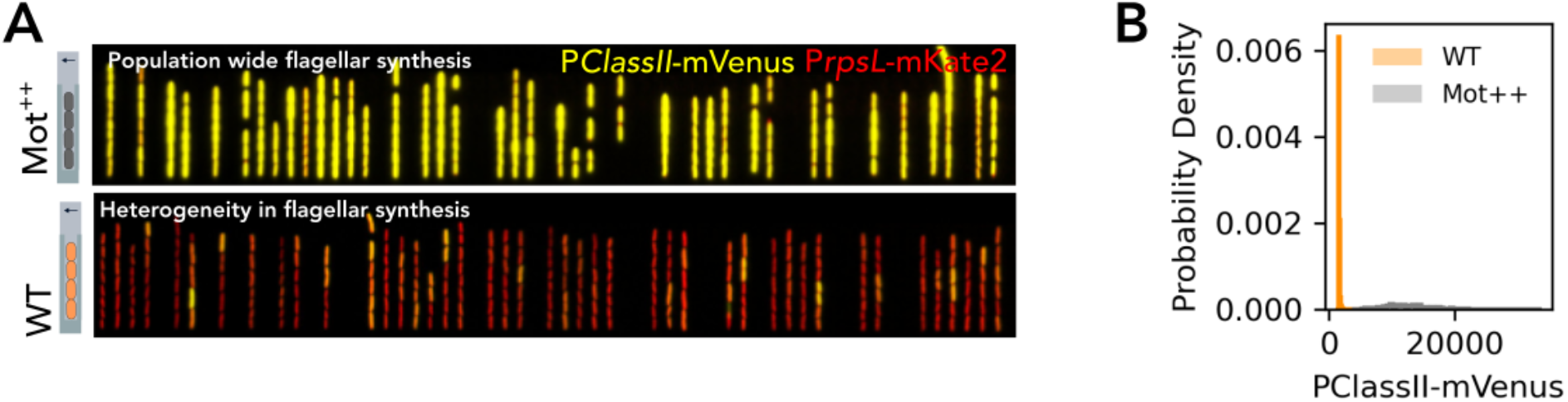
Mot++ constitutively expresses flagellar cascade. (A) Representative kymographs comparing Class II gene expression (PClassII-mVenus, yellow) overlaid with a constitutive marker (PrpsL-mKate2, red) in hyper-motile (Mot++, top) versus WT (bottom) cells in conditioned media. (B) Probability density distribution of PClassII-mVenus expression, demonstrating constitutive high activity in Mot++ (grey, CV=0.5) compared to the low, heterogeneous activity in WT (orange, CV=1.0).

**Figure S5.**
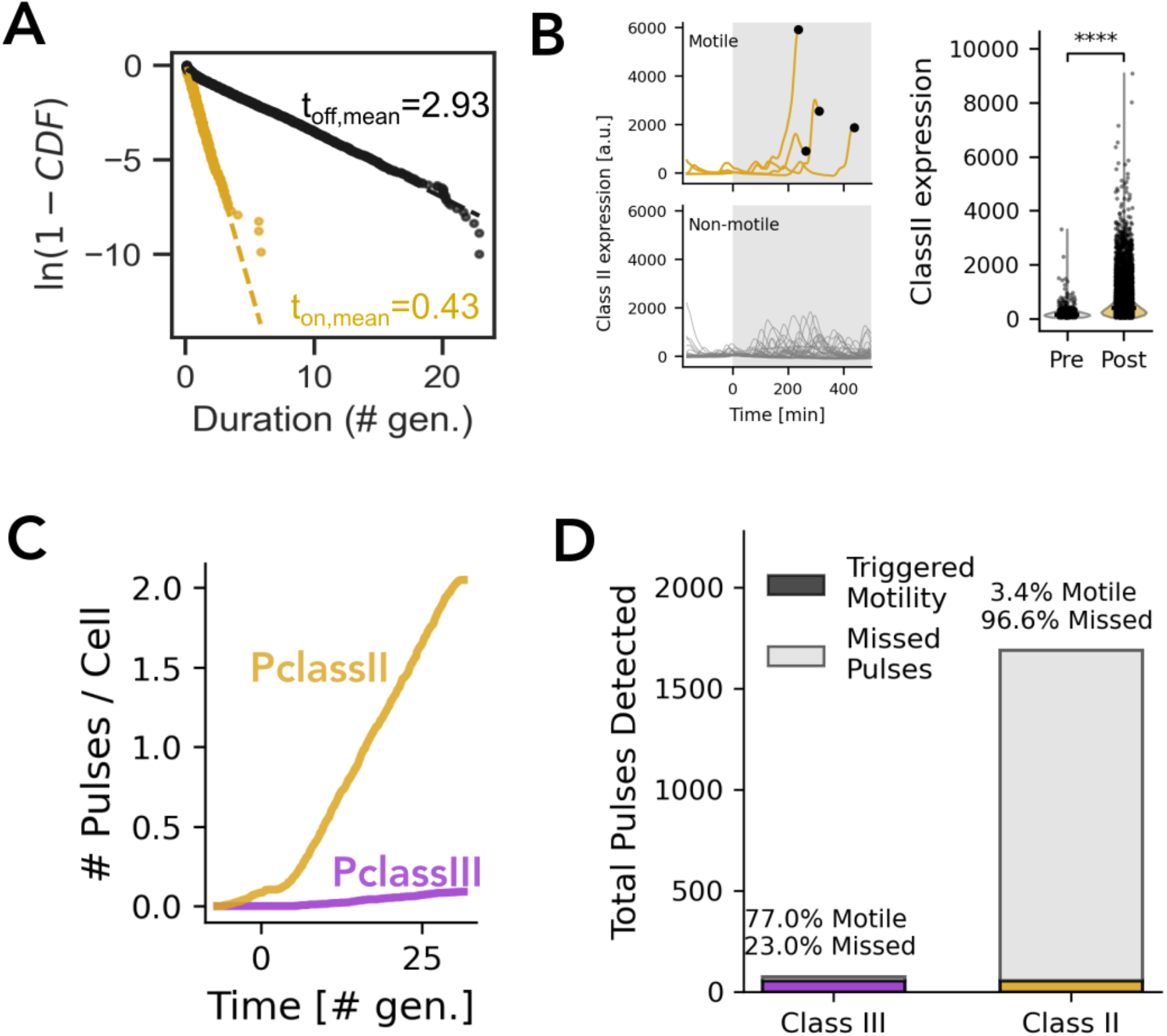
Most Class II pulses are not followed by motility, whereas Class III activation closely precedes motility in WT. (A) Cumulative distribution function plotted as ln(1-CDF) versus duration in generations for P*classII* promoter activity (on: yellow, off: black). (B) (left) Single-cell traces of Class II expression in motile (top) and non-motile (bottom, grey) mother cells. The grey shaded region denotes the switch to conditioned media at t=0 mins. Black dots mark the moment of cell exit (motility onset). (right) Distribution of Class II peak intensity for cells growing in fresh media (pre) exposed to conditioned media (post). (C) Cumulative number of Class II (Yellow) and Class III (Purple) pulses per cell over time for WT cells with conditioned media provided at t=0 mins. (D) Quantification of the single-reporter pulse success in the bar chart with fraction of missed pulses in grey and pulses that triggered motility in colored (Class III in purple, and Class II in yellow).

**Figure S6.**
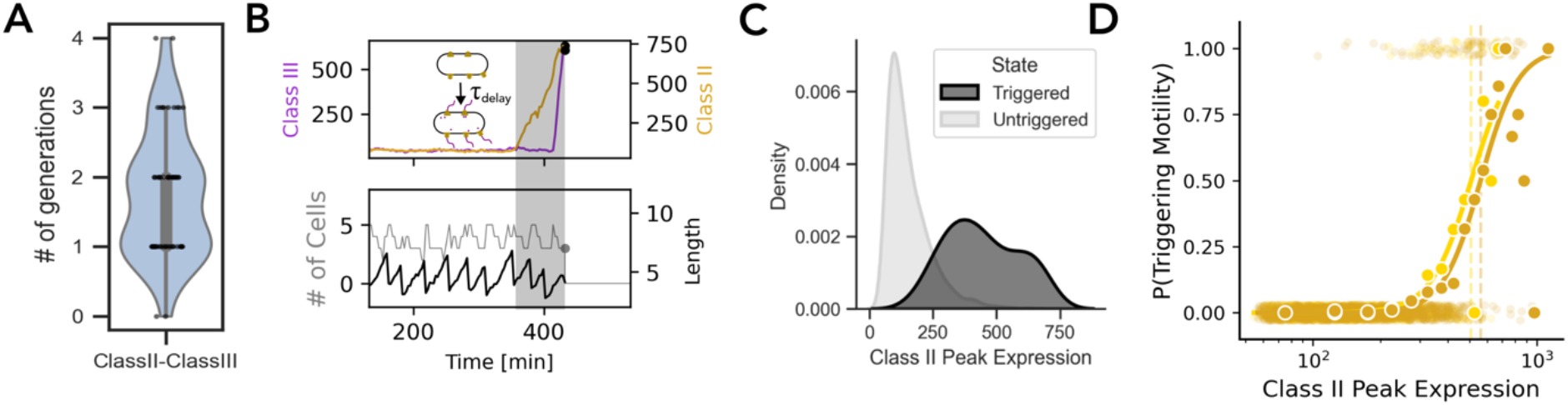
Commitment to motility is gated by a threshold of Class II expression and is coordinated among lineage related cells. (A) Violin plots quantifying the delay in number of generations from the activation of Class II to activation of Class III for motile cells. (B) Representative single-cell traces tracking Class II and Class III expression (top) alongside cell length and division events (bottom) leading up to cell exit. (C) Corresponding probability density distributions for peak P*classII* expression amplitude between motility triggering (dark grey) and pulses not followed by motility (light grey). (D) Probability of triggering motility plotted against peak Class II expression for 2 repeats of experiments. Faint scatter points represent binary single-cell outcomes (0 = untriggered, 1 = triggered), while solid yellow dots represent binned probabilities. The solid yellow line indicates a Hill equation fit. Vertical dashed lines denote the expression corresponding to the 50% commitment threshold.

**Figure S7.**
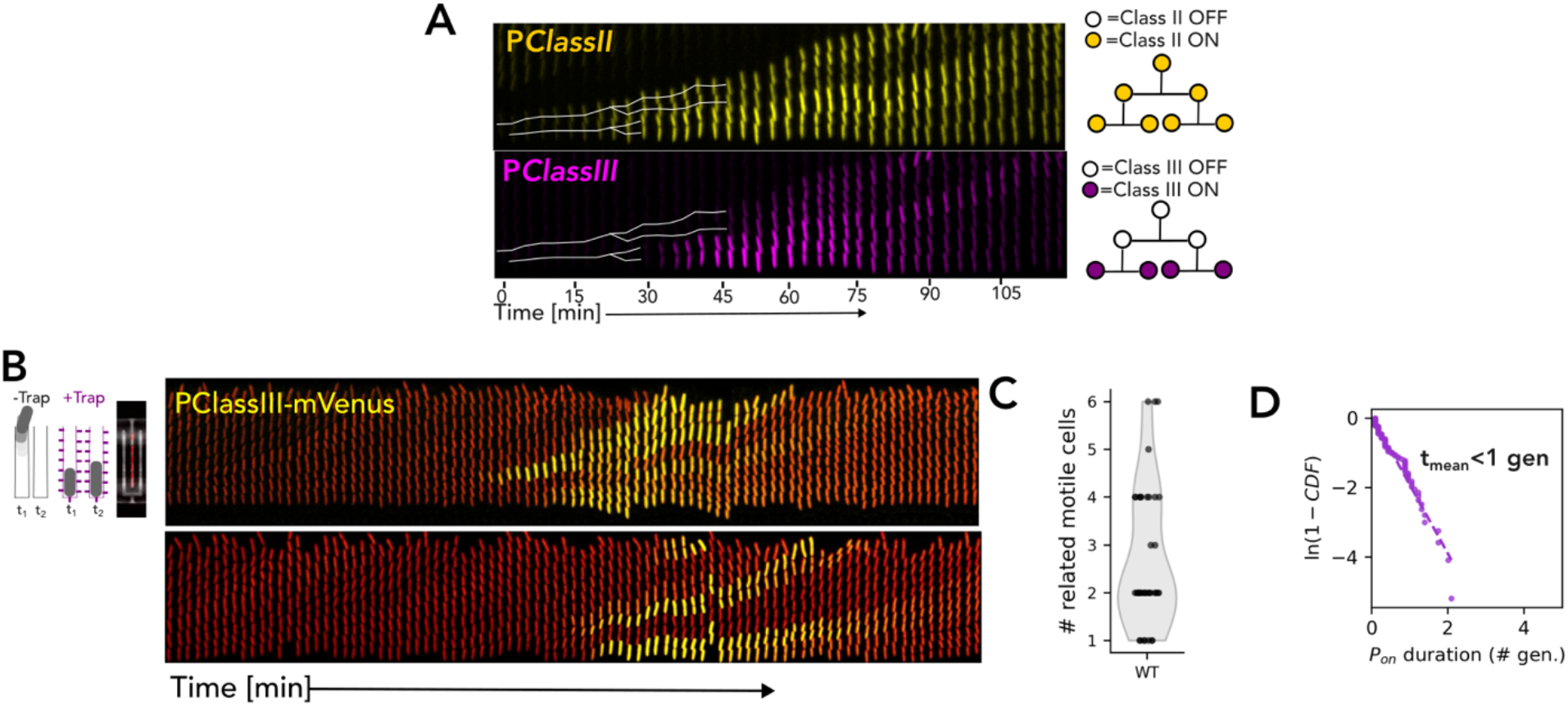
Class III activation is coordinated among related cells retained in the trapping device. (A) (left) Representative kymographs from the trap device showing (top) P*ClassII* and (bottom) P*ClassIII* activity. White lines designate multi-generational lineages as they appear in kymographs. (right) Lineage tree tracing the activation of Class II and Class III for kymograph depicted on the left. (B) (Left) Schematic comparing the conventional microfluidic mother-machine (-Trap) with a modified design utilizing back-pressure to physically retain swimming cells (+Trap). (Right) Kymographs showing *PClassIII-mVenus* activity (yellow) overlaid on constitutive *PrpsL-mKate2* (red) for growth channel with a motile mother cell in trap chip. (C) Violin plot quantifying the number of related cells within a lineage that coordinate activation of Class III expression. (D) Cumulative distribution function (plotted as ln(1−CDF)) of the Pon duration (in generations) for Class III expression.

**Figure S8.**
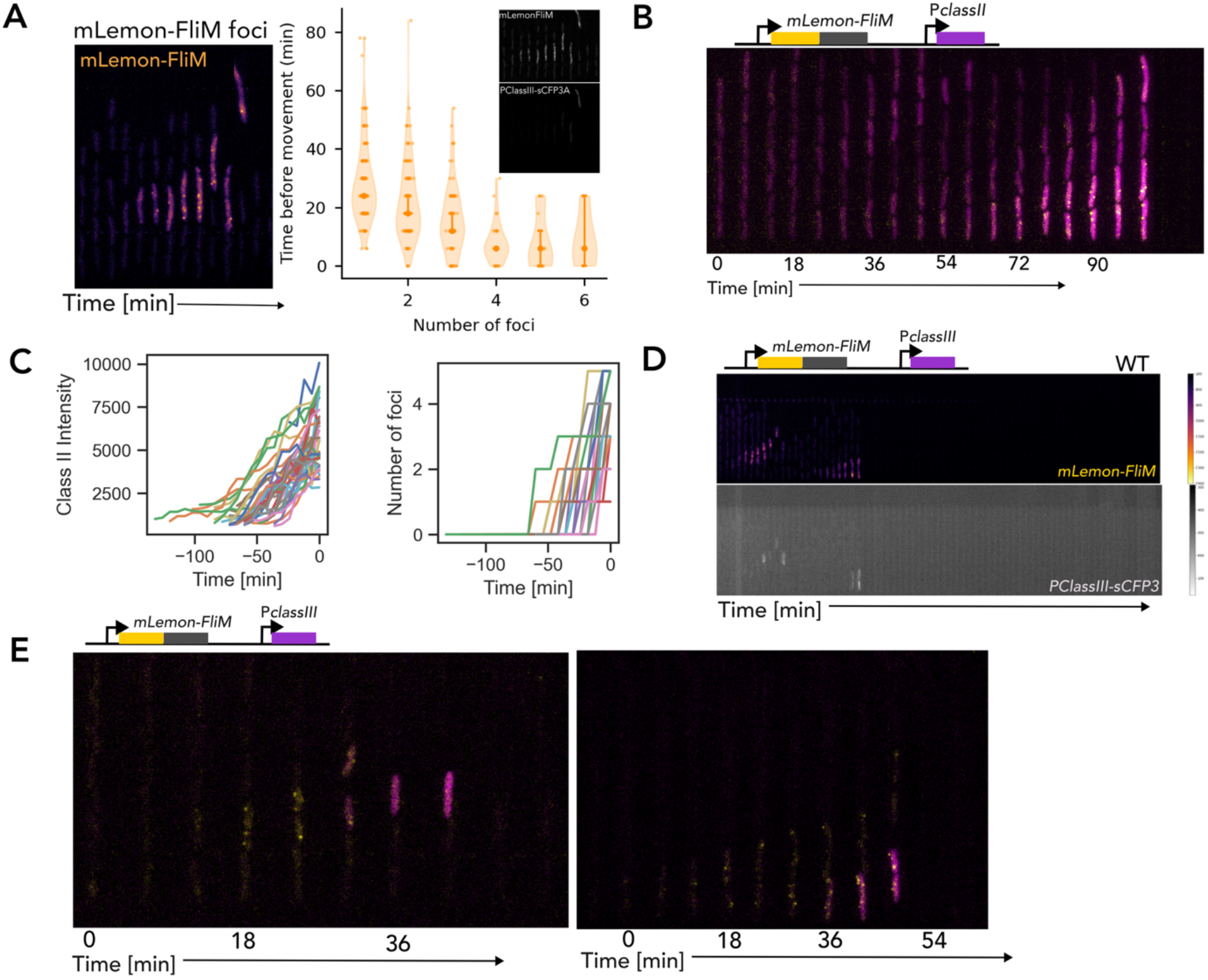
Structural accumulation of basal bodies dictates the timing of motility commitment. (A) (Left) Kymograph capturing the progressive accumulation of basal bodies (*mLemon-FliM* foci, orange) within a cell lineage prior to motility onset. (Right) Violin plots quantifying the temporal delay to movement as a function of the number of FliM foci that a motile lineage started with. Cells one to three foci exhibit longer and more variable delays, whereas cells with more than three foci become motile sooner. Inset shows *mLemon-FliM* foci (top) over time along with *PClassII-sCFP3* (bottom). (B) (Top) Schematic of the dual-reporter construct. (Bottom) Kymograph tracking basal body assembly (*mLemon-FliM*, yellow) alongside Class II transcription (*PClassII-sCFP3*, magenta) simultaneously within the same lineage. (C) Single-cell mother cell trajectories of (Left) Class II transcriptional intensity and (Right) the accumulation of FliM foci, aligned to the exact moment of physical cell exit (t = 0). (D) (Top) Schematic of the dual-reporter construct. (Middle and Bottom) Kymographs tracking basal body assembly (*mLemon-FliM*, top) alongside Class III transcription (*PClassIII-sCFP3*, bottom, grayscale) simultaneously within the same lineage. (E) (Top) Schematic of the dual-reporter construct. (bottom) Kymographs tracking basal body assembly (*mLemon-FliM*, yellow) alongside Class III transcription (*PClassIII-sCFP3*, magenta) simultaneously within the same lineage.

**Figure S9.**
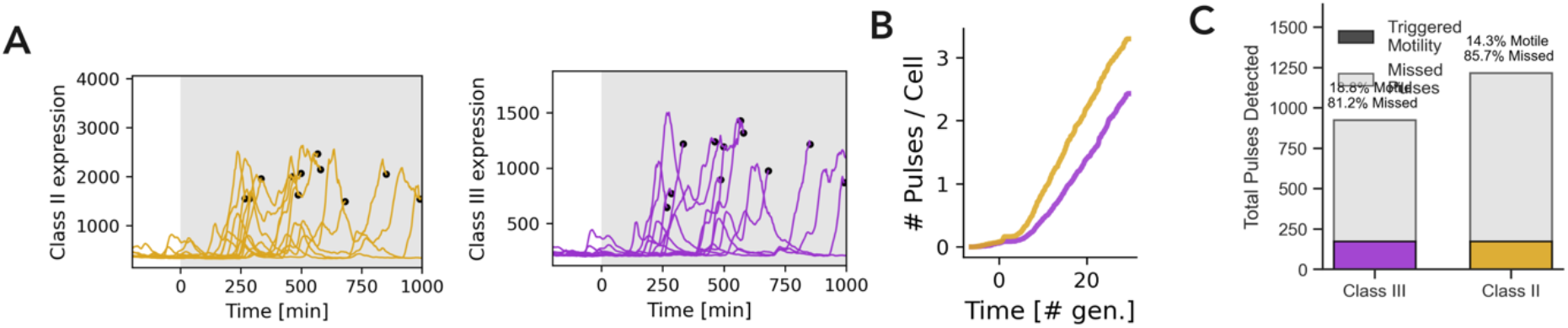
Characterization of the motility decision architecture and transcriptional commitment for *ΔflgM* cells. (A) Single-cell traces of (Left, yellow) Class II and (Right, purple) Class III expression in motile *ΔflgM* mother cells. The grey shaded region denotes the switch to conditioned media. Black dots mark the moment of cell exit (motility onset). (B) Cumulative number of Class II (Yellow) and Class III (Purple) pulses per cell over time for *ΔflgM* cells with conditioned media provided at t=0 mins. (C) Quantification of the single-reporter pulse success in the bar chart with fraction of missed pulses in grey and pulses that triggered motility in colored (Class III in purple, and Class II in yellow) for *ΔflgM* cells.

**Figure S10:**
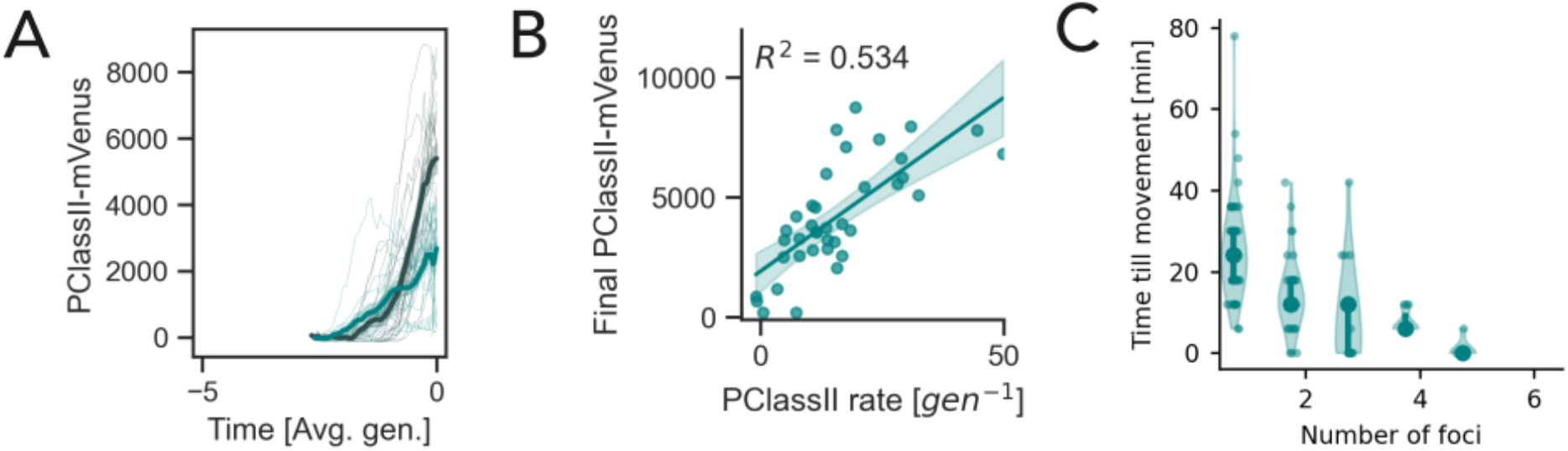
Class II expression kinetics and basal body accumulation of motile *ΔflgM* lineage. (A) Class II gene expression traces for *ΔflgM* cells comparing fast (black) versus slow (teal) lineages aligned to the onset of motility (t = 0). (B) Scatter plot showing the correlation between the final peak Class II expression and the rate of Class II accumulation per generation in *ΔflgM* lineages shown in Panel A. (C) Violin plots quantifying the temporal delay to movement as a function of the number of FliM foci in *ΔflgM*.

## Supplementary Movies

**Movie 1.** Time lapse of Class II flagellar promoter expression (P*classII*-mVenus, top) and P*rpsL*-mKate2 (bottom) for *E. coli* Mot++ cells growing in microfluidic trenches, sequentially washed with media filtered from a batch culture sampled every 30 minutes across its growth curve. Scale bar = 5 *μm*.

**Movie 2.** Time lapse of Class III flagellar promoter expression (P*classIII*-mVenus, yellow) overlaid onto constitutive expression of P*rpsL*-mKate2 (red) for *E. coli* cells growing in microfluidic trenches under conditioned media treatment introduced at t=0 mins. Scale bar = 10 *μm*.

**Movie 3.** Time lapse of Class II flagellar promoter expression (P*classII*-mVenus, yellow) overlaid onto constitutive expression of P*rpsL*-mKate2 (red) for *E. coli* cells growing in microfluidic trenches under conditioned media treatment introduced at t=0 mins. Scale bar = 10 *μm*.

**Movie 4.** Time lapse of P*rpsL*-mKate2 (top), Class III flagellar promoter expression (P*classIII*-mVenus, middle) and Class II flagellar promoter expression (P*classII*-sCFP3A, bottom) for *E. coli* cells growing in microfluidic trenches with traps under conditioned media treatment introduced at t=0 mins. Scale bar = 10 *μm*.

**Movie 5.** Time lapse of mLemon-FliM reporter (yellow foci) in *E. coli* cells growing in microfluidic trenches with traps under conditioned media treatment. Scale bar = 10 *μm*.

**Movie 6.** Time lapse of *PClassII-sCFP3A* (top) and *mLemonFliM* (bottom) for *E. coli* cells growing in microfluidic trenches with traps under conditioned media. Scale bar = 5 *μm*.

**Movie 7.** Time lapse of *mLemonFliM* (top) and *PClassII-sCFP3A* (bottom) for *E. coli* cells growing in microfluidic trenches with traps under conditioned media. Scale bar = 5 *μm*.

**Movie 8.** Time lapse of *mLemonFliM* (top) and *PClassIII-sCFP3A* (bottom) for *E. coli* cells growing in microfluidic trenches under conditioned media. Scale bar = 5 *μm*.

**Movie 9.** Time lapse of P*rpsL*-mKate2 (top), Class III flagellar promoter expression (P*classIII*-mVenus, middle) and Class II flagellar promoter expression (P*classII*-sCFP3A, bottom) for Δ*flgM E. coli* cells growing in microfluidic trenches with traps under conditioned media treatment introduced at t=0 mins. Scale bar = 5 *μm*.

**Movie 10.** Time lapse of mLemon-FliM reporter expression (top) and P*classIII*-sCFP3A expression (bottom) for Δ*flgM E. coli* cells growing in microfluidic trenches with traps under conditioned media treatment. Scale bar = 10 *μm*.

## Materials and Methods

### Strains, Plasmids, and Genetics

All experiments were performed using the *Escherichia coli* K-12 strain MG1655 as the wild-type background. To enable robust lineage tracking and morphological segmentation during microfluidic cultivation, the strain was engineered to constitutively express the cytoplasmic fluorescent marker *PrpsLmKate2*. Chromosomal deletion mutants (Δ*fliC*, Δ*flgM*) were generated via P1 phage transduction using donor strains from the Keio collection^62^. Following transduction, the kanamycin resistance cassettes were excised using the Flp recombinase expression vector pCP20 to generate markerless deletions. To monitor the temporal dynamics of the flagellar hierarchy at single-cell resolution, transcriptional reporters were constructed using regulatory regions corresponding to the Class II and Class III stages. The Class II promoter (*PrpoF* or *PClassII*) was derived from the upstream region of *fliA* (bp −90 to −1 relative to the transcription start site). Crucially, this precise 90-bp fragment was selected because it encompasses the complete dual-promoter architecture governing fliA expression. It captures the intact 46-bp FlhD₄C₂ binding (GTAACCCCCAAATAACCCCTCATTTCACCCACTAATCGTCCGATTA) required for the initial σ^70^-dependent activation of the cascade. Furthermore, the sequence natively retains the downstream −35 and −10 core promoter elements (ATTAAAAACCCTGCAGAAACGGATAAT) specific to the secondary, FliA (σ^28^)-dependent promoter. By incorporating these overlapping binding sites, the *PrpoF* reporter preserves the full dual-input logic of the flagellar switch: primary activation by the master regulator complex and the subsequent positive autoregulatory feedback loop mediated by FliA^10,47^. For the Class III reporter, an 85-bp promoter region of *flgM* (*PflgM* or *PClassIII*) was utilized. This fragment was designed to encompass the FliA-dependent canonical binding elements, capturing both the −35 (TAAAGATT) and −10 (GCCGATAA) core promoter motifs. In its native context, *flgM* is transcribed both as part of the Class II *flgAMN* operon and from an internal FliA-dependent promoter located immediately upstream of *flgM*. The 85-bp fragment cloned here spans only the latter: the FlhD4C2-dependent promoter driving *flgAMN* lies upstream of flgA, outside the cloned region, so the reporter is read out from the σ^28^ promoter alone and reports free FliA rather than FlhD4C2 activity^10^. The dual-reporter plasmids were engineered using a modular Golden Gate assembly framework^63^ and integrated into a low-copy plasmid backbone (pSC101 origin). The synthetic parts—comprising promoters, ribosome binding sites (RBS), fluorescent reporters, and terminators—were synthesized with flanking BsaI recognition sequences (‘GGTCTC’) and amplified with AGGCACTTGCTCGTACGACG and ATGTGGGCCCGGCACCTTAA overhangs. The promoters were coupled to the spectrally distinct fluorescent reporters mVenus and sCFP3a, respectively^64^. A medium strength Weiss Ribosome Binding Site synthetic (lib197, a MoClo basic part modified from Bba_B0032 to adjust spacing) was inserted immediately upstream of both start codons to ensure consistent translation initiation and high-sensitivity detection. To eliminate transcriptional interference and read-through, the two reporter modules were separated by strong Voigt_DT3^65^ double transcriptional terminators and oriented in tandem. The final constructs were assembled in a single-pot restriction-ligation reaction utilizing BsaI and T4 DNA Ligase. Strict directional assembly and correct ordering of the reporter modules were enforced via custom junction sequences, which generated precise 4-basepair overhangs to link the components into the finalized pSC101 backbone. All assembled plasmids were verified via whole plasmid sequencing with Plasmidsaurus. To track the accumulation of HBB by tagging the flagellar C-ring under its native regulation, an N-terminal translational fusion of the fluorescent reporter mLemon to FliM was engineered. Because fliM is naturally transcribed as the second gene within the fliLMN operon, expressing it in isolation can disrupt native translational coupling and expression stoichiometry. To prevent this, the construct was designed to encompass the complete native fliLMN operon promoter along with the full coding sequence of the upstream gene, fliL, which lies upstream of fliM in the native operon. The endogenous intergenic spacing between the fliL stop codon (TAA) and the fliM start codon was strictly preserved to maintain physiological ribosome binding dynamics. The mLemon coding sequence was integrated at the native fliM translation start site. To physically decouple the folding of the fluorophore from the structural assembly of the FliM rotor protein, the C-terminus of mLemon was fused to FliM via a flexible, genetically encoded hexaglycine linker^66^ (GGAGGCGGAGGCGGAGGCG). To maintain the continuous open reading frame and avoid steric clashes at the N-terminus, the linker was joined to the third amino acid of native FliM (Aspartate, GAT), effectively replacing its native Start-Methionine and subsequent Glycine. The complete PfliL-fliL-mLemon-linker-fliM-DT3 cassette was assembled into the low-copy pSC101 backbone using the standardized Golden Gate protocol.

### Media and Growth Conditions

Strain construction was performed in LB broth or LB supplemented with antibiotics (25 *μ*g/mL chloramphenicol) at 37°C. 4 mL cultures were grown at 220 rpm in 20mL glass culture tubes. Successfully constructed strains were stored in glycerol stocks at −80°C. For experiments, strains were streaked from these glycerol stocks on LB agarose plates with appropriate antibiotic selection. A single colony was picked and then grown overnight in growth media. For all microfluidic experiments, cells were grown in EZ Rich Defined Medium (EZRDM) supplemented with 0.2% glucose. Conditioned medium was prepared by filtering a 200 mL bulk culture of MG1655 (1/50 overnight dilution) at specific times during the growth phases. This filtered medium was delivered to the microfluidic device via peristaltic pump at a constant flow rate of 240 *μ*L/min. To establish conditions that induce motility and to validate channel exit as a behavioral readout, we collect cell-free medium from batch cultures at 30-min intervals across the growth curve and sequentially flush it through the device **[Figure 1B, S1]**. Medium collected at an OD of approximately 0.7 produces the largest motile fraction in Mot++ and is therefore used as conditioned medium.

### Soft agar assay

For motility assays on soft agar, 1 *μ*L of overnight culture was spotted on freshly prepared and autoclaved 0.2% tryptone agar plates. The cultures were incubated at 37°C for 16 hours before capturing images. Mot++ strain was isolated from the outer ring of the swarm assay. We tested for the presence of insertion in the promoter of *flhDC* using GCCATGCATTACAGAACATCG and GTAAAGACCCATTTCTATTTG, and confirmed the sequence by whole genome sequencing using Plasmidsaurus.

### Microfluidic Device Fabrication

Single-cell imaging was performed using the ‘mother machine’ microfluidic device^40^. The chip has a main channel for flow of media. This main channel is branching into perpendicular growth channels (or called growth trenches). Two distinct microfluidic designs were utilized: a standard mother-machine and a specialized trapping chip. Mother Machine: Growth trenches (25 X 1.4 X 1.4 *μ*m) were oriented perpendicular to a main flow channel (25 X 100 *μ*m) Trapping Chip: This design incorporated additional nanometer sized side-ports spanning the growth trenches, connected to a back-side flow channel. This architecture provided sufficient backpressure to physically retain motile cells while allowing constant media exchange. The chips were made of polydimethylsiloxane (PDMS) polymer using a silicon wafer mold. A 1:10 solution of polymerising agent and PDMS monomer were rigorously mixed and then poured onto the silicon wafer. This was placed in a vacuum chamber to remove air bubbles. The device was then heated at 95°C in an oven for 2 h to polymerize. For each experiment, one chip was cut out using a scalpel, and holes for inlet and outlet were inserted using a 0.7 mm biopsy puncher. The PDMS chip was bonded on a glass coverslip (thickness No 1.5). The cleaned coverslip and PDMS chip were exposed to air plasma for 2 min and bonded at 95°C for 30 min.

### Mother machine setup

100 μL of an overnight culture was introduced in the microfluidic chips by pipetting through the inlet. The chip was then inserted into a custom-built centrifuge holder and spun at 650 rpm for 1.5 min to aid the loading of cells into the growth trenches and then reversed to load the other side of growth trenches with spinning at 400 rpm for 30 seconds. Glass bottles with caps allowing for tubing inlets and pressurization were filled with EZRDM media or spent media as indicated. The silicon tubing (Tygon) was loaded onto peristaltic pump to deliver media into chips at a constant flow rate.

### Live-Cell Imaging and Microscopy

Time lapse imaging was performed using a Nikon Ti-2 inverted fluorescence microscope equipped with Plan Apo λ 40x Ph2 DM air objective or Plan Apo λ 60x Oil objective (for visualizing FliM foci), Prior motorized stage, Iris15 camera, laser excitation source (Celesta), and operated with a perfect focus system. Exposure times were 300 ms for mKate2 (λ = 555 nm), 80 ms for CFP reporters (λ = 440 nm) and 200 ms for mVenus (λ = 508 nm) using 100% of maximal LED excitation intensities. The excitation and emission lights were separated using a triband dichroic and individual emission filters. The microscope chamber (Okolabs) was maintained at 37°C throughout the experiments. Images were captured every 3 min or 6 minutes (for FliM foci experiments).

### Dilution model

To ask how the kinetics of a Class II pulse set the timing of motility, we simulated a minimal model in which transcriptional output is converted into flagellar structures. The model is dimensionless; time is measured in generations and expression in arbitrary units.

A cell activates Class II expression at a constant rate *r* for a duration *τ*, after which transcription stops, and the product is diluted by growth:

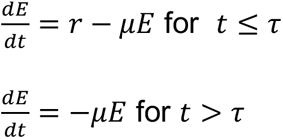

Giving:

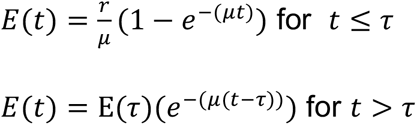

Solving for peak amplitude for t= *τ*, we get:

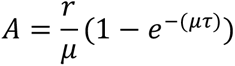

Here *μ* =ln2 per generation, so dilution alone halves the Class II pool once per division. Division is represented as continuous dilution of the lineage rather than as discrete partitioning between daughters. Because *τ* is held constant across cells, A is proportional to *r*, and cell-to-cell variation in pulse size arises entirely from variation in activation rate.

Class II expression generates a pool of unassembled precursor, or burden, *B*. This pool is consumed by an assembly machinery of limited capacity and simultaneously diluted by growth:

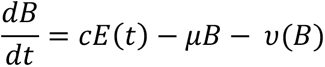

where *c* is the precursor produced per unit Class II expression and *υ* (B) is the assembly rate, capped at a maximum *υmax* and limited by the available burden, *υ* (B) = min (*υmax*, B/Δt), with B constrained to be non-negative. Because assembly saturates, a larger pulse deposits a burden that takes proportionally longer to clear, so the model predicts that the delay to motility grows with the size of the pulse. A cell is scored as committing to motility once the burden has been built up and subsequently cleared: we record the first time at which B < 0.02 following the first time at which B > 0.2, tolerances that stand in for B = 0 and avoid the asymptotic approach to zero inherent to the numerical solution.

Equations were integrated by the forward Euler method with a time step Δt = 0.02 generations over 25 generations, for 100 independent cells with activation rates *r* drawn from a lognormal distribution (mean 0.5, standard deviation 0.4 of the underlying normal) and a pulse duration τ held constant across cells. Simulations used a fixed random seed. Parameter values are *c*= 1.0, *υmax* = 0.8 and *τ* = 2.5 generations.

### Image Processing and Lineage Tracking

Time lapse microscopy data were saved as nd2 files and visualized in Fiji^67^. The data were processed using the BACMMAN^68^ plugin in Fiji and further analyzed using custom Python scripts. Plots and analysis were generated using Python. Images were first pre-processed by BACMMAN using the *PrpsLmKate2* fluorescence channel to stack all individual growth trenches and correct for experimental drift in x-y coordinates and image rotation. The outlines of cells in the growth trenches were then jointly segmented and tracked over time based on the *PrpsLmKate2* fluorescence signal. The traces were visually inspected and manually corrected for errors in segmentation or lineage tracing using the BACMMAN software. Reporter fluorescence was extracted by overlaying the cell masks from the *PrpsLmKate2* channel onto the CFP and mVenus channels and computing the mean intensity over the cell area. BACMMAN generated output in excel files containing cell growth characteristic. *PrpsLmKate2* intensity data, other fluorescence intensity data. These files were then further analyzed using a custom python pipeline.

### Quantification of Population Motile Fractions

To quantify the population-wide motility kinetics, the cumulative motile fraction (fmotile) was calculated by tracking the physical retention of mother cells located at the closed end of the microfluidic growth trenches. For each analyzed microfluidic position, the initial number of tracked mother cells was recorded at the beginning of the experiment (N0). At each subsequent imaging frame (t), the number of remaining mother cells was counted (Nt). The cumulative fraction of motile cells that had successfully exited the trench was computed as fmotile(t)=1−(Nt/N0).

To account for inter-experiment variability, single-cell binary exit events were pooled across multiple independent biological replicates and distinct microfluidic channel positions for each strain evaluated (Wild-type, *ΔflgM*, Mot++, and *ΔfliC*). Imaging frames were converted to absolute physical time (hours) and coordinated such that t=0 corresponds to the exact moment conditioned media was introduced into the main flow channel.

Due to the asynchronous nature of single-cell exits, the pooled longitudinal data for each strain was discretely binned into standardized 0.1-hour (6-minute) temporal windows. Within each bin, the mean motile fraction and the standard error of the mean (SEM) were calculated. On resulting population kinetic plots, solid lines represent the binned mean fmotile, while the shaded bounding regions denote the ± SEM to indicate variability in the estimated mean.

### Channel exit definition for mother cells and tracking cell dynamics pre exit

Channel exit or a motile cell was defined when all cells in a growth channel were displaced. To quantify gene expression dynamics leading up to motility, single-cell tracking data from the microfluidic trenches were processed using custom Python scripts. The analysis was performed for the bottom-most “mother” cell of each trench (index = 0) and its direct lineage. To ensure data quality and exclude morphological anomalies (such as filamentation), cell tracks were filtered to include only those with a minimum trajectory of 5 frames, a maximum cell length of less than 13 μm. Fluorescence intensities for the dual reporters (e.g., Class II and Class III promoter activity) were quantified over time. For temporal alignment and visualization, the frame numbers were converted to physical time (minutes) relative to the time of the media switch. Single-cell trajectories of fluorescence expression, cell length, and the total number of cells residing in the corresponding trench were plotted aligned to the exact time of cell exit. The visualization window was standardized to display the dynamics from a fixed interval prior to cell exit until after exit, clearly capturing the drop in trench cell count to zero as the motile cell washed out.

### Signal Processing and Transcriptional Peak Classification

To distinguish stochastic transcriptional fluctuations from pulses associated with subsequent motility, raw fluorescence intensities for the dual reporters (Class II and Class III) were subjected to signal processing. Single-cell fluorescence trajectories were first smoothed using a centered 5-frame moving average to minimize frame-to-frame imaging noise. To remove a per-trace intensity offset, each trace was baseline-corrected by subtracting the background fluorescence.

Algorithmic peak detection (via the SciPy find_peaks module in Python) was then applied to the smoothed, baseline-corrected traces. Transcriptional peaks were identified requiring a minimum temporal separation of 5 frames and a defined prominence threshold tailored to the specific reporter fluorophore to ensure only genuine transcriptional bursts were captured.

Detected peaks were subsequently classified into two functional categories: ‘missed’ (futile) pulses and ‘triggered’ (committal) pulses, based on the physical fate of the cell: Missed Pulses: For lineages that remained physically retained in the growth channel until the end of the experiment, all detected transcriptional peaks were defined as non-committal, missed pulses.

Triggered Pulses: For lineages that successfully transitioned to motility (indicated by premature track truncation), the terminal fluorescence measurement immediately preceding channel exit was designated as the triggered motility peak. Any independent transcriptional peaks occurring earlier in these same exiting lineages were classified as missed pulses.

### Growth, Elongation, and Dilution-Corrected Promoter Activity

Interdivision times were extracted from continuous single-cell lineages during steady-state pre-treatment conditions to calculate the mean generation time (Td). The specific growth rate (μ) for the cellular population was subsequently defined as μ=ln(2)/Td. To capture dynamic morphological changes, the instantaneous single-cell elongation rate was computed as the temporal derivative of the natural logarithm of cell length, d(lnL)/dt. To quantify transcriptional dynamics independent of passive cell growth and fluorophore dilution, Promoter Activity (PA) was calculated from the smoothed fluorescence intensity (I). We computed PA by taking the numerical temporal derivative of the fluorescence signal and adding a growth-dilution correction term, defined as PA= dI/dt+μI.

### Population Motility Kinetics and Bulk Culture Alignment

To directly correlate single-cell motility decisions within the microfluidic device to macroscopic population growth phases, microfluidic tracking data was temporally aligned with bulk culture optical density (OD) measurements. The cumulative fraction of motile cells in the microfluidic device (f_motile_) was quantified by tracking the occupancy of individual growth channels over time. This metric captures motility events as the physical exit of the mother cell from the trench. To align this with bulk physiology, independent batch cultures were grown under identical media conditions, and their OD was recorded at discrete intervals. These bulk OD time points were mapped onto the continuous microfluidic time axis, perfectly synchronized to the moment of the conditioned media switch. Simultaneously, mean population-level fluorescence trajectories for Class II and Class III reporters were generated by averaging the normalized single-cell intensities across all tracked lineages at each corresponding frame, allowing for direct temporal comparison between population-level growth limits, average transcriptional shifts, and the physical onset of swimming behavior.

### Statistical Analysis and Mathematical Modeling of the Sensitive Commitment Threshold

To quantify the function governing the transition to motility, single-cell outcomes were assessed as a probability function of their peak Class II expression amplitude. The binary fate of triggering motility for each pulse (Y∈{0,1} for Untriggered vs. Triggered) was mapped against pulse magnitudes of P*classII* expression. This unbinned binary data was fitted to a non-linear Hill-Langmuir probability function: 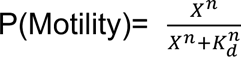 where P(Motility) represents the probability of a transcriptional pulse culminating in physical cell exit, X is the peak Class II expression amplitude, Kd is the half-maximal activation threshold, and n is the apparent Hill coefficient.

The parameters n and Kd were extracted using non-linear least-squares optimization (scipy.optimize.curve_fit), bounded to physiologically relevant limits. A higher n signifies a steeper, more digital commitment boundary, whereas the Kd value defines the expression corresponding to a fitted motility probability of 0.5.

### Quantification of *mLemonFliM* foci

To simultaneously monitor transcriptional activation and physical motor assembly at single-cell resolution, lineages were tracked using a dual-reporter system combining a Class II or Class III transcriptional reporter (e.g., *PClassIIsCFP3* or *PClassIIIsCFP3*) with a hook-basal body (HBB) tag (*mLemonFliM*). Microfluidic image data was processed using the BACMMAN software. For transcriptional quantification, cells were automatically segmented, and the mean whole-cell fluorescence intensity was extracted for each tracked lineage. Because the transition to motility is a rare phenotypic event and structural basal bodies manifest as exceptionally dim, diffraction-limited puncta, structural quantification required a targeted, hybrid approach. Microfluidic trenches exhibiting successful motility events were visually identified, cropped, and converted into longitudinal single-cell kymographs using BACMMAN. The *mLemonFliM* foci were quantified manually from these kymographs. Individual foci were validated by extracting spatial line scans across the longitudinal axis of the cell using Fiji or NIS-Elements software, checking that the localized peak intensity exceeded the local cytoplasmic baseline fluorescence. To quantify the temporal coupling between Class II transcription and *mLemonFliM* foci appearance, single-cell tracking data was processed using custom Python scripts (utilizing Pandas and SciPy libraries). Raw transcriptional traces and discrete foci counts were temporally aligned. To isolate the active phase of transcription, the cells were followed from the point of exit to where the gene expression was above the baseline expression. Within this active window, the rate of Class II reporter accumulation and the rate of increase in visible FliM foci were calculated for each individual cell using linear regression (scipy.stats.linregress). The lineages were sorted based on their specific Class II accumulation rates. The expression trajectories and foci accumulation curves of the fastest and slowest assembling lineages were aggregated to generate mean dynamic traces (± SEM), allowing for direct comparison of structural integration windows across phenotypic extremes.

### Lineage level coordination of motility

To visualize the spatial and temporal coordination of flagellar gene expression across entire related lineages, the single-cell tracking analysis was expanded beyond the isolated mother cell (Idx = 0) to encompass all residing descendants within a given microfluidic growth trench. For lineages that successfully executed a motility event (validated by mother cell exit), the spatial coordinate along the longitudinal axis of the trench was extracted for every cell.

To map transcriptional states to physical lineage positions, single-cell fluorescence intensities for both Class II and Class III reporters were mapped to these spatial coordinates over time. This multidimensional array was projected as a quantitative spatial kymograph, allowing for the direct visual assessment of how transcriptional bursts spatially propagate through sister and cousin cells occupying the same trench prior to their collective exit. Lineage coordination was defined as the commitment to motility among related descendants sharing a microfluidic trench. The degree of multi-generational coordination was mathematically quantified as log2(N), where N represents the absolute number of related cells participating in a coordinated motility burst. This logarithmic metric describes group size on a division-equivalent scale; it does not by itself measure elapsed generations or establish genealogical relationships.

The coordination was quantified using strain-specific physical and transcriptional proxies: Wild-Type (WT) Lineages: In WT cells, Class III transcription functions as a proxy for complete basal-body structural assembly and imminent motility. Thus, lineage coordination was directly quantified by measuring the number of related cells within a trench exhibiting simultaneous Class III transcriptional bursts immediately preceding the coordinated washout event. *ΔflgM* Lineages: In *ΔflgM* mutants, the premature release of FliA uncouples Class III expression from actual structural completion, rendering Class III bursts invalid as a proxy for physical motility. Therefore, coordination in the mutant was quantified by measuring the extended temporal integration window itself. Specifically, calculating the number of cellular generations elapsed between the initial activation of Class II reporter and the actual physical onset of movement (trench exit).

---

**Table S1.**
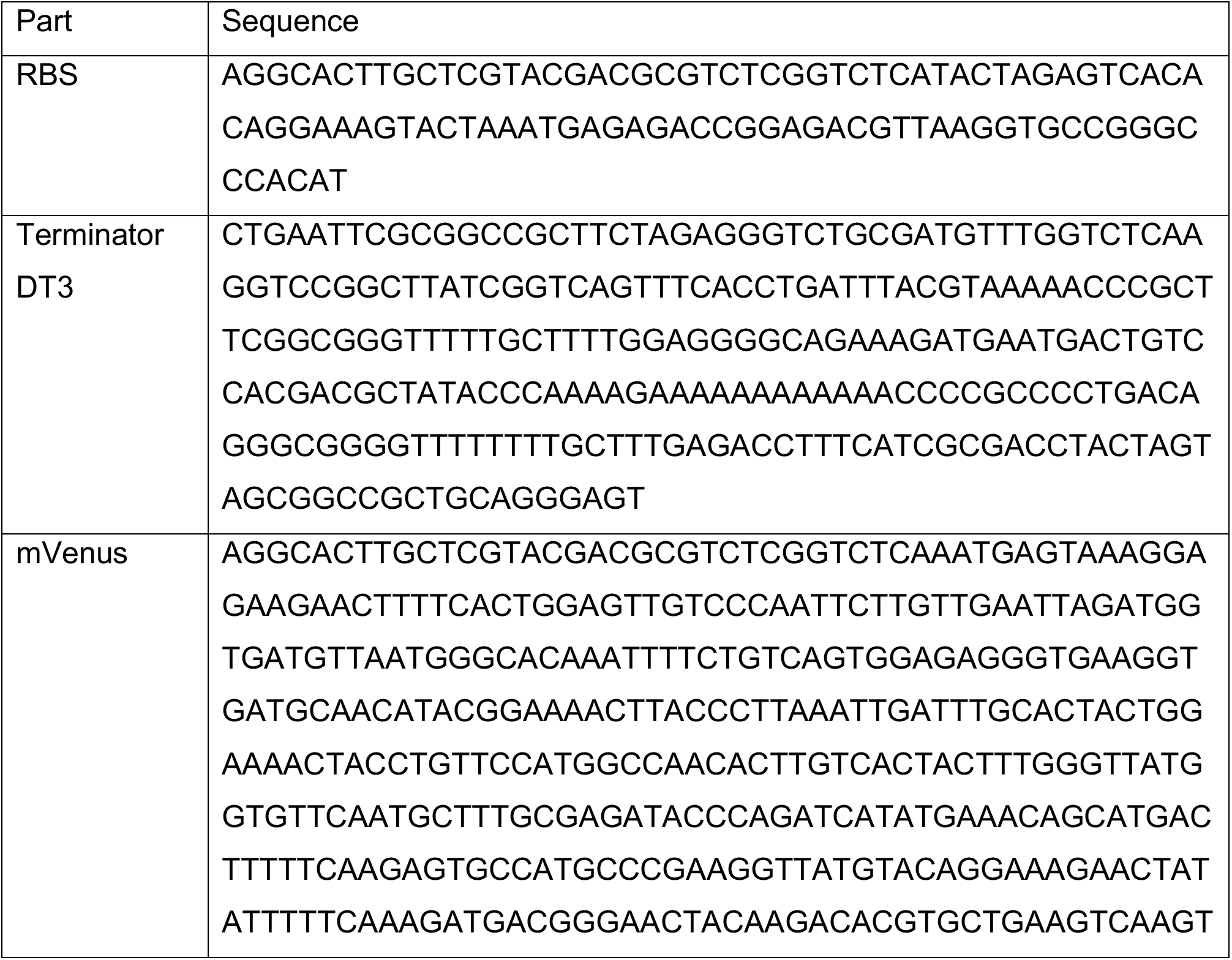
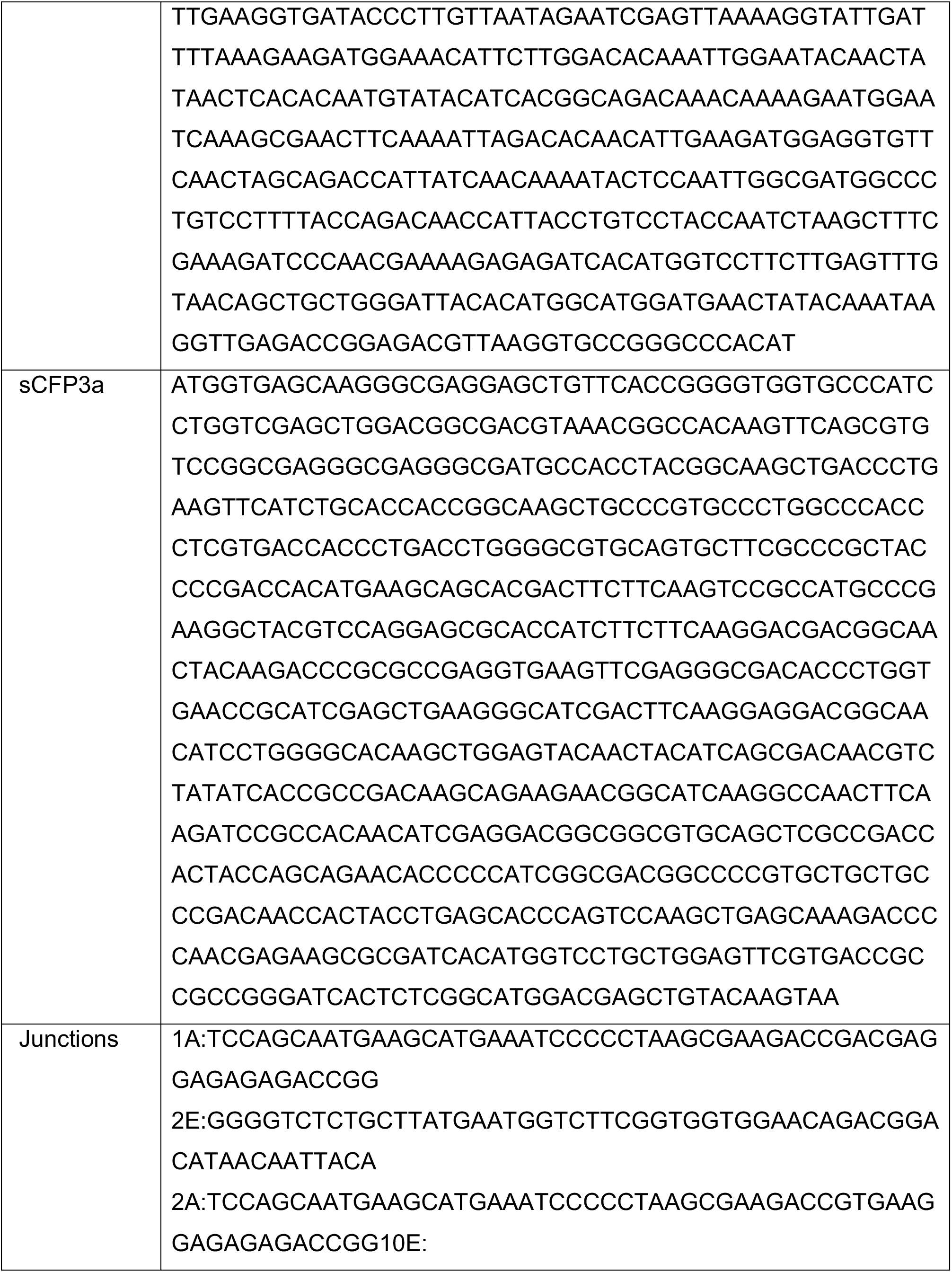
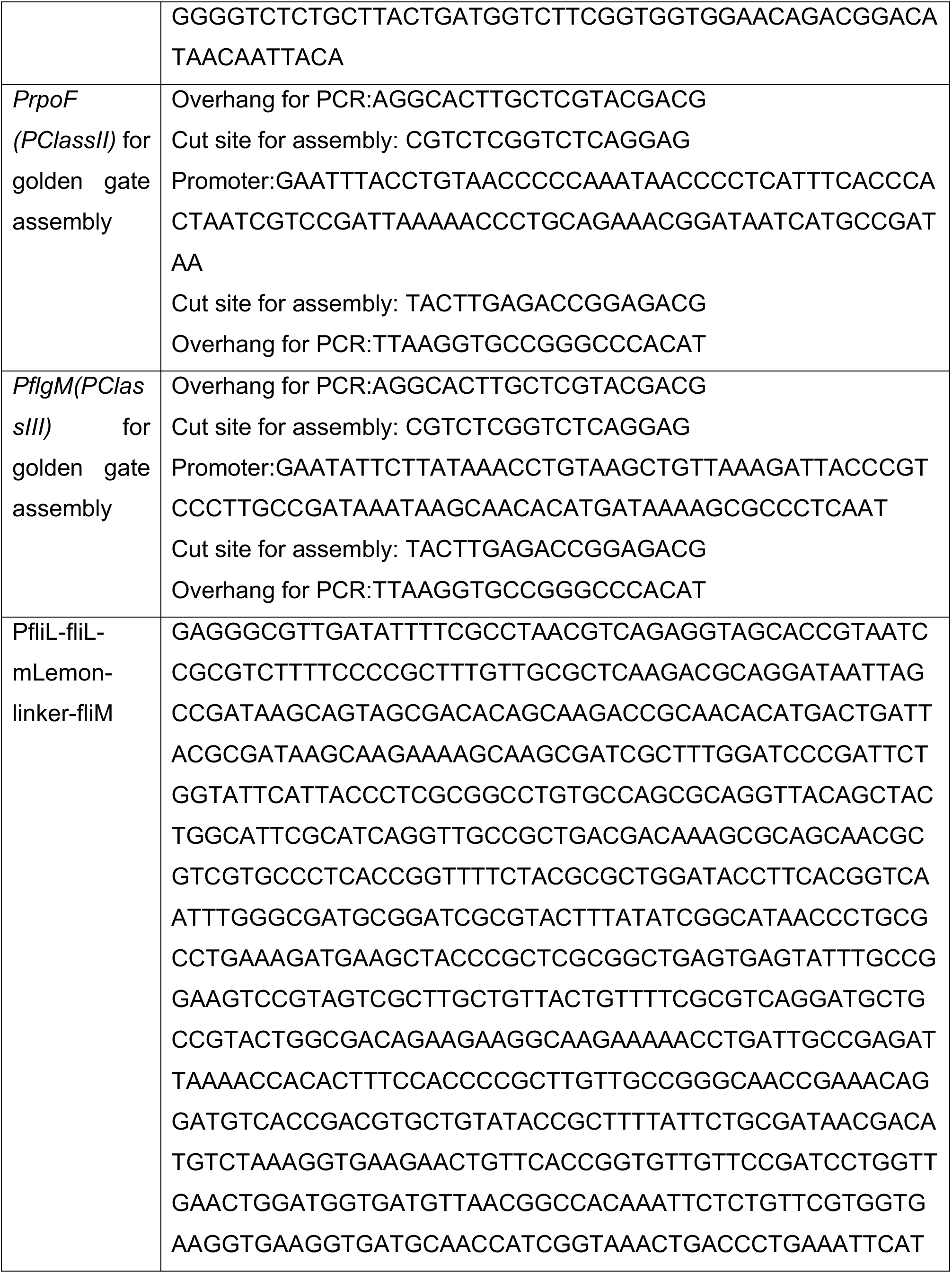
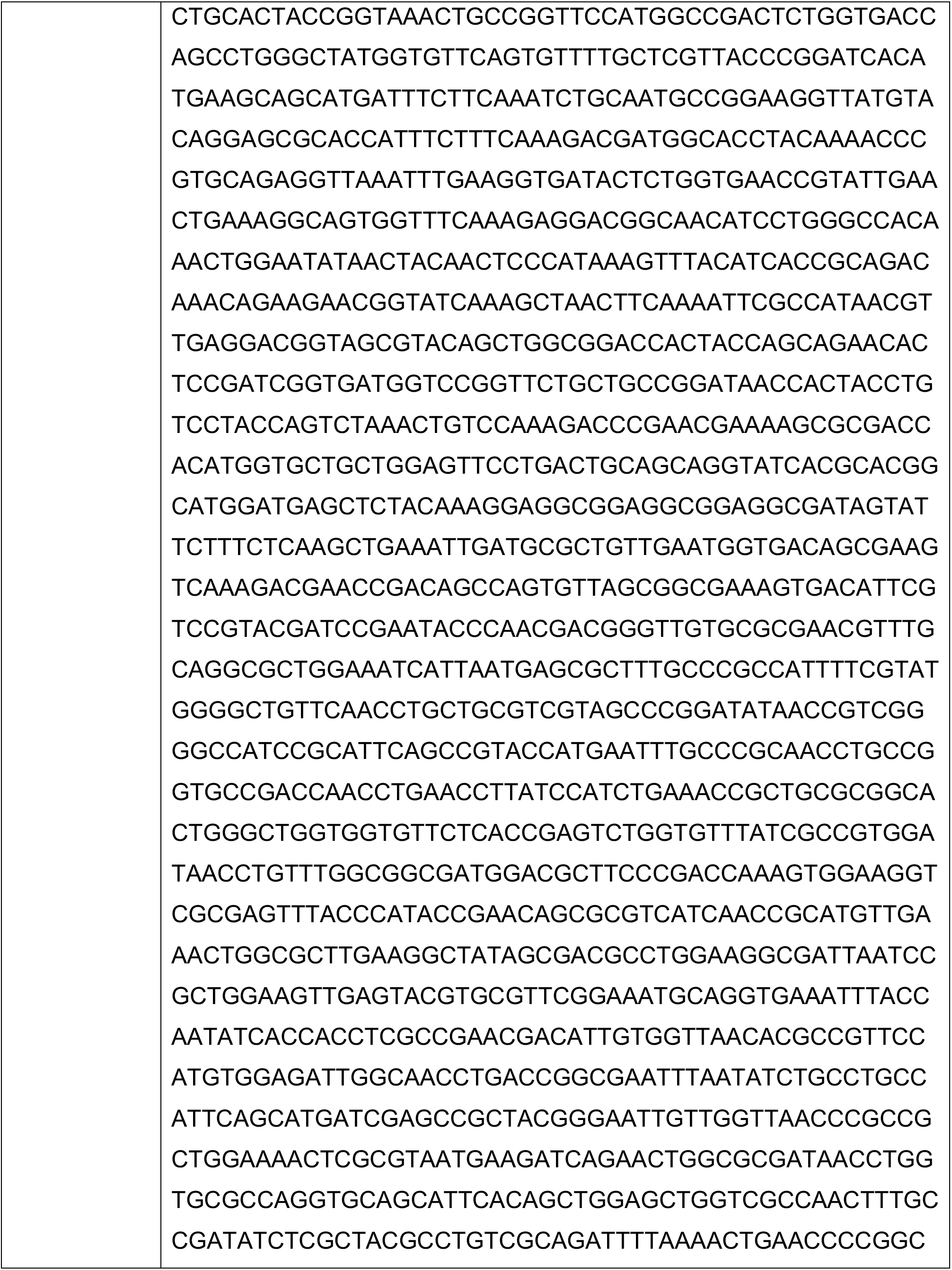
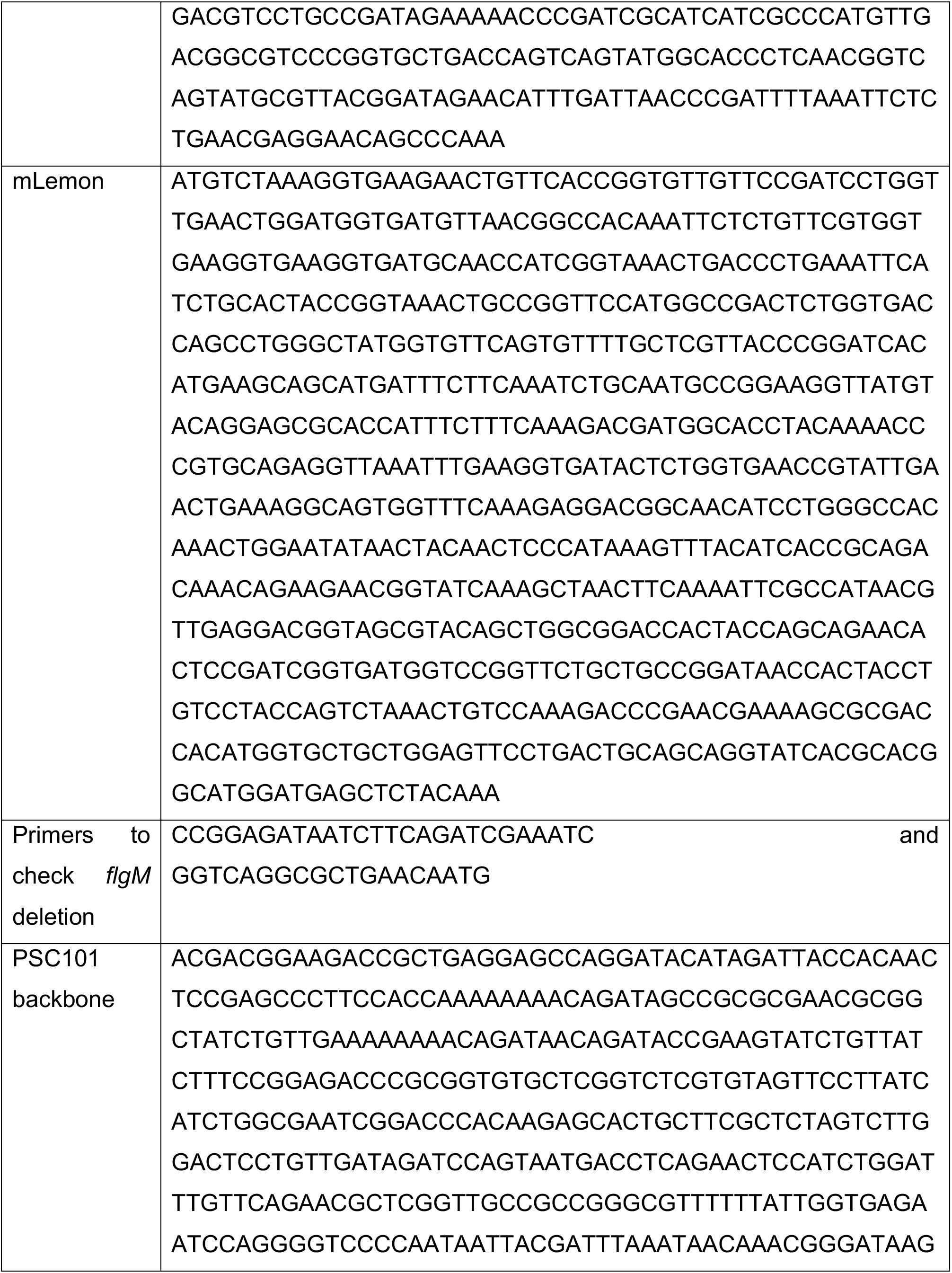
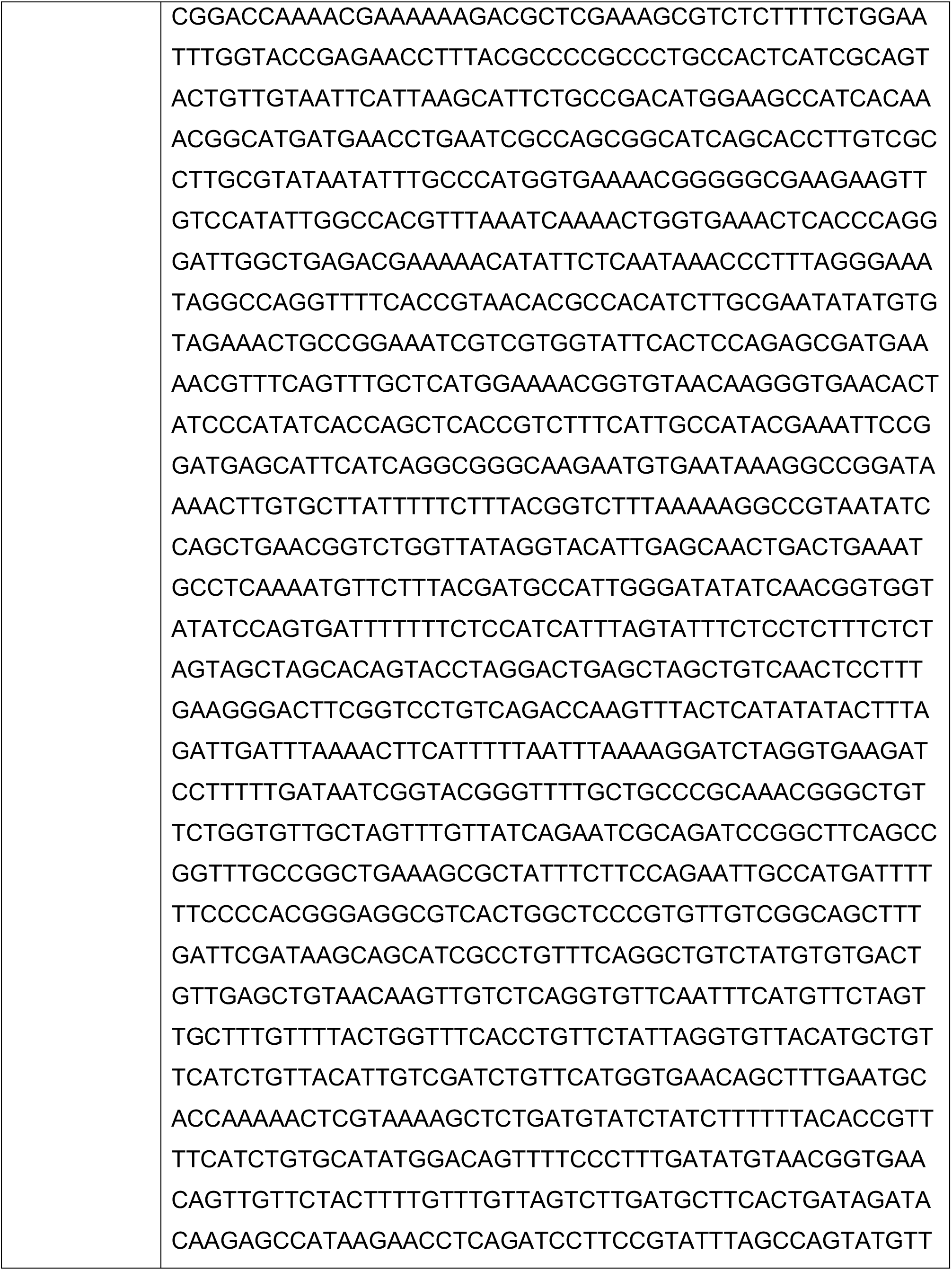
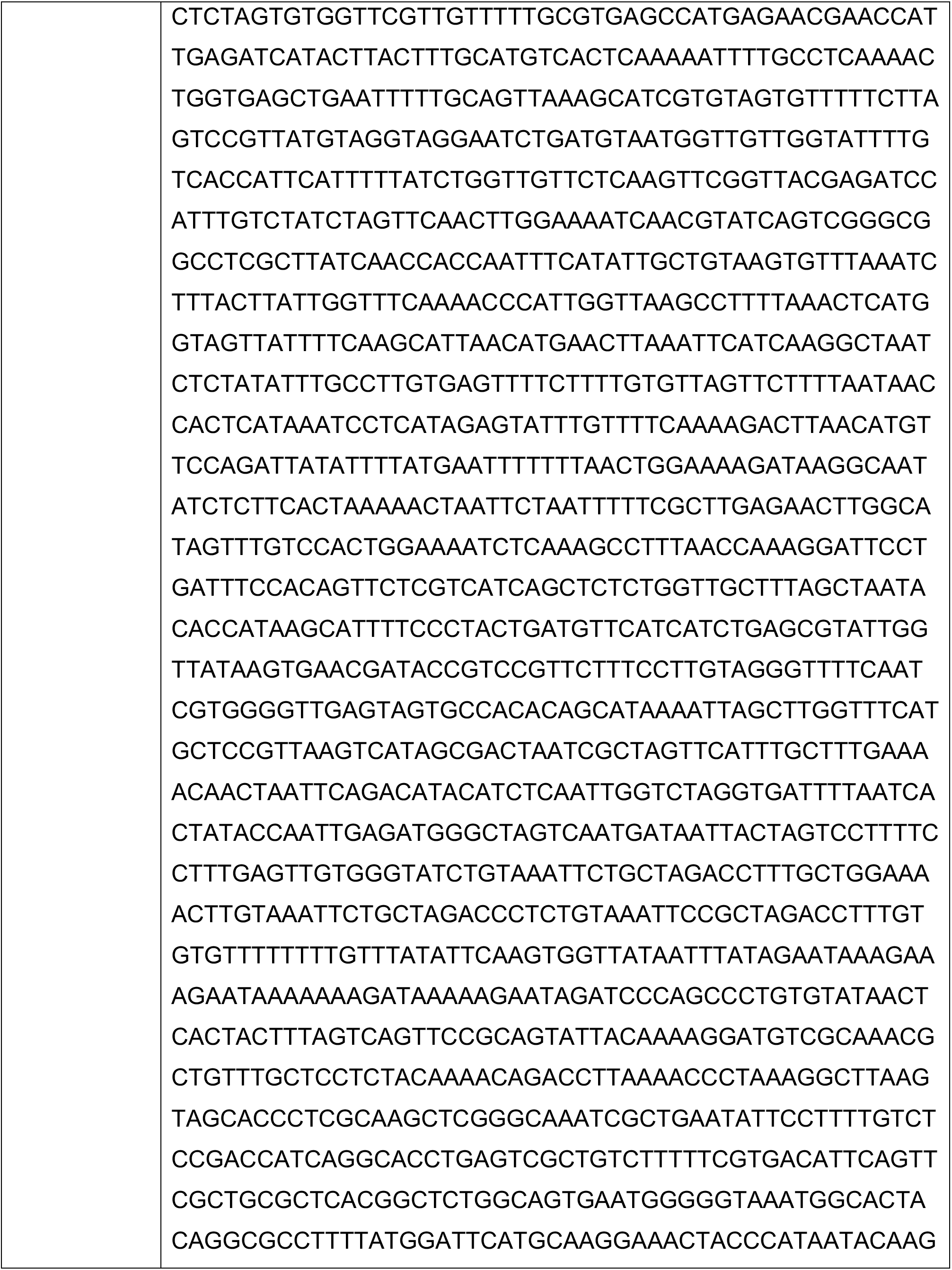
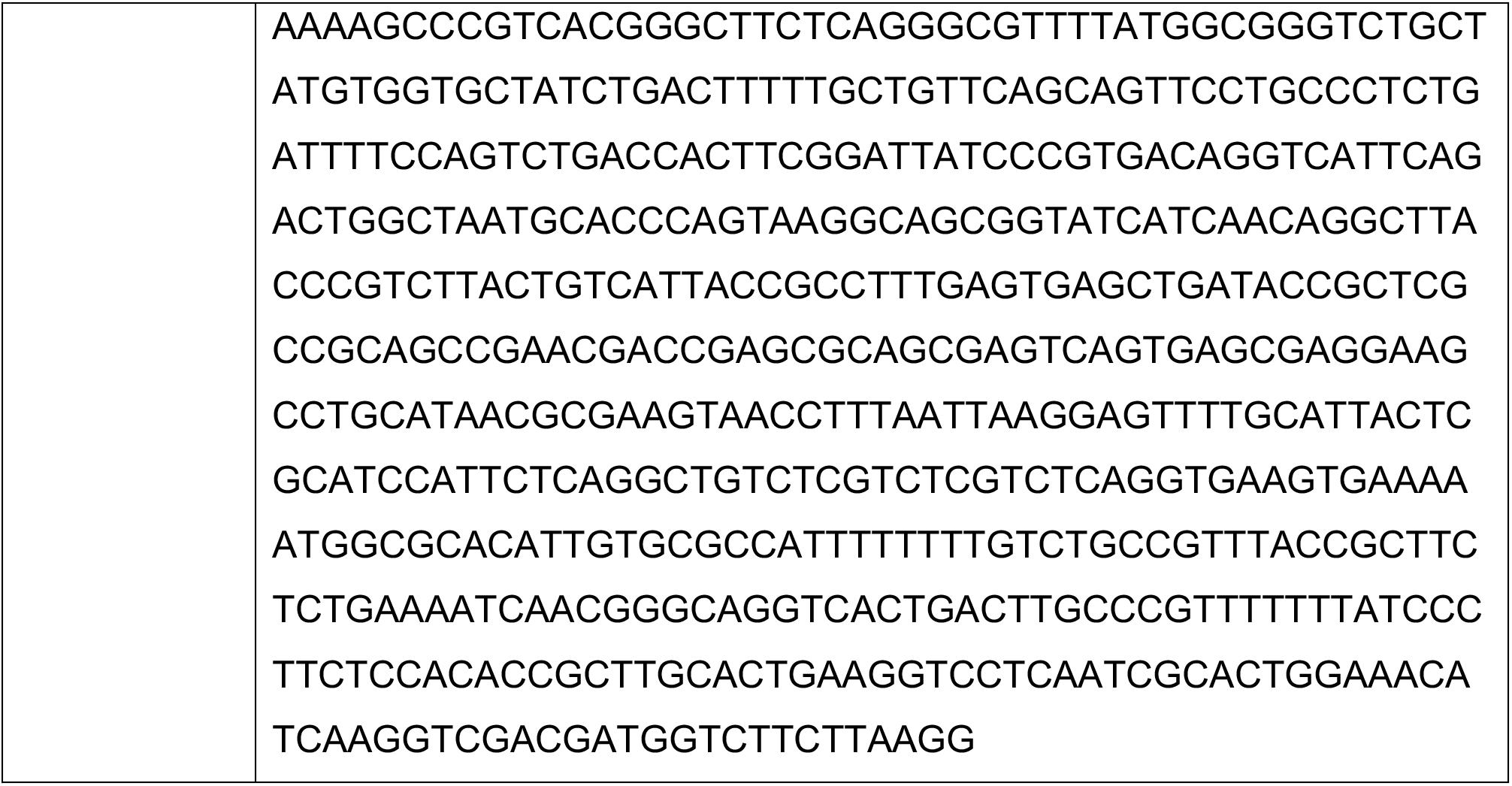
DNA parts and sequences.

